# Limited population structure but signals of recent selection in introduced African Fig Fly *(Zaprionus indianus*) in North America

**DOI:** 10.1101/2024.09.20.614190

**Authors:** Priscilla A. Erickson, Alexandra Stellwagen, Alyssa Bangerter, Ansleigh Gunter, Nikolaos T. Polizos, Alan O. Bergland

## Abstract

Invasive species have devastating consequences for human health, food security, and the environment. Many invasive species adapt to new ecological niches following invasion, but little is known about the early steps of adaptation. Here we examine population genomics of a recently introduced drosophilid in North America, the African Fig Fly, *Zaprionus indianus*. This species is likely intolerant of subfreezing temperatures and recolonizes temperate environments yearly. We generated a new chromosome-level genome assembly for *Z. indianus*. Using resequencing of over 200 North American individuals collected over four years in temperate Virginia, plus a single collection from subtropical Florida, we tested for signatures of population structure and adaptation within invasive populations. We show founding populations are sometimes small and contain close genetic relatives, yet temporal population structure and differentiation of populations is mostly absent across North America. However, we identify two haplotypes that are differentiated between African and invasive populations and show signatures of selective sweeps. Both haplotypes contain genes in the cytochrome P450 pathway, indicating these sweeps may be related to pesticide resistance. X chromosome evolution in invasive populations is strikingly different from the autosomes, and a haplotype on the X chromosome that is highly differentiated between Virginia and Florida populations is a candidate for temperate adaptation. These results show that despite limited population structure, populations may rapidly evolve genetic differences early in an invasion. Further uncovering how these genomic regions influence invasive potential and success in new environments will advance our understanding of how organisms evolve in changing environments.

**Article Summary:** Invasive species (organisms that have been moved outside their natural range by human activities) can cause problems for both humans and the environment. We studied the genomes of over 200 individuals of a newly invasive fruit fly in North America, the African Fig Fly. We found genetic evidence that these recently introduced flies may be evolving in their new environments, which could make them stronger competitors and more likely to become pests.

## Introduction

Understanding how species expand and adapt to new environments in an era of changing land use, environmental changes, and global commerce is central to controlling the spread of disease (Altizer et al. 2013; Hoberg and Brooks 2015), to maintaining crop security (Oerke 2006; Sutherst et al. 2011) and to preserving biodiversity (Bellard et al. 2012). Many organisms are moving to new, previously unoccupied ranges at rates that continue to accelerate (Ricciardi 2007; Seebens et al. 2015; Seebens et al. 2017; Platts et al. 2019; Sardain et al. 2019) due to changing environmental conditions, habitat alteration, and anthropogenic introductions. Genetic adaptation to new environments may allow some vulnerable organisms to survive in new habitats but may also permit potentially harmful organisms to expand even further (Clements and Ditommaso 2011). The past two decades have produced a wealth of studies characterizing the genetic and genomic basis of adaptation in a variety of organisms, from experimental populations of microbes (Good et al. 2017; Nguyen Ba et al. 2019; Johnson et al. 2021) to natural populations of eukaryotes (Hancock et al. 2011; Jones et al. 2018; Barrett et al. 2019; Lovell et al. 2021 Jan 27; Schluter et al. 2021). Recent and ongoing invasions offer the opportunity to study rapid evolution and adaptation to new environments in nearly real-time (Koch et al. 2020; Pélissié et al. 2022; Parvizi et al. 2023; Soudi et al. 2023). Recently, genomics has helped trace the history and sources of many well-known invasions (Pélissié et al. 2022; Picq et al. 2023) and shown that genetic divergence and even local adaptation are common in invasive populations that have been established for decades or even centuries (Ma et al. 2020; Stuart et al. 2021; Li et al. 2023). However, much remains unknown about the role that evolution plays in allowing invasive organisms to colonize and thrive in new environments. A better understanding of adaptive pathways in invasion may assist in predicting the success of invasions and controlling their outcomes.

The African Fig Fly, *Zaprionus indianus*, serves as a unique model to study how invasion history and local environment influence patterns of genetic variation. The ongoing, recurrent invasion of *Z. indianus* in North America offers a premier opportunity to study the possibility of rapid genetic changes following invasion. The *Zaprionus* genus arose in Africa but Z. *indianus* was first described in India in 1970 (Gupta 1970), where it has adapted to a range of environments (da Mata et al. 2010). It is one of the most ecologically diverse drosophilids in Africa; its ability to utilize up to 80 different food sources (Yassin and David 2010) and its generation time of as few as ∼13 days (Nava et al. 2007) likely fueled its spread around the world. In 1999, it was first detected in Brazil (Vilela 1999), where it subsequently spread and caused major damage to fig and berry crops as well as native fruit species (Leão and Tldon 2004; Oliveira et al. 2013; Roque et al. 2017; Zanuncio-Junior et al. 2018; Allori Stazzonelli et al. 2023). It was later found in Mexico and Central America in 2002-2003 (Markow et al. 2014) and eventually Florida, USA in 2005 (Linde et al. 2006). In 2011-2012, its range expanded northwards in eastern North America (Joshi et al. 2014; Timmeren and Isaacs 2014; Pfeiffer et al. 2019) and eventually reached as far north as Ontario, Canada (Renkema et al. 2013) and Minnesota, USA (Holle et al. 2018). It has also recently been found in the Middle East, Europe, and Hawaii (Parchami-Araghi et al. 2015; Kremmer et al. 2017; Willbrand et al. 2018), suggesting that the invasion is ongoing. *Z. indianus* can damage fig and berry crops (Pfeiffer et al. 2019; Allori Stazzonelli et al. 2023), increasing concerns about its pest potential in its expanding range.

Despite its global success, *Z. indianus* males are sterile below 15 °C, making cold temperatures a limiting factor to their success (Araripe et al. 2004). Within the temperate environment of Virginia, the species exhibits strong seasonal fluctuations in abundance (Rakes et al. 2023). First detection in Virginia is usually in June or July, weeks after the appearance of other overwintering drosophilids, and population sizes climb dramatically through the late summer and early fall, when it often dominates the drosophilid community in temperate orchards. Typically, the peak in early to mid-September is followed by a dip in abundance and then a second peak in October, suggesting a seasonal component to reproduction or fluctuations in factors influencing *Z. indianus’* relative fitness. However, despite its early post-colonization success, it does not appear to survive temperate winters; *Z. indianus* populations became undetectable in Virginia by early December (Rakes et al. 2023). *Z. indianus* were not collected during regular winter sampling in Memphis, Tennessee (Kohlmeier and Kohlmeier 2025), and they were collected in lower numbers during the winter in northern Florida relative to central Florida (Renkema et al. 2018), suggesting that even mild winters (only 8 nights below 0° C) can substantially reduce their numbers. In locations in Minnesota, Kansas, and the northeastern US, *Z. indianus* has been detected one year and then not the next, suggesting that the populations may be extirpated by cold and re-introduced by stochastic dispersal processes (Holle et al. 2018; Gleason et al. 2019; Rakes et al. 2023). While conclusive data are still lacking, these results collectively suggest *Z. indianus* likely repeatedly invades temperate environments each year from warmer refugia. Reinvasion offers an opportunity to recurrently study the genetic impacts of invasion and the potential for post-colonization adaptation across multiple years of sampling.

Genetic *s*tudies *of Z. indianus* are limited but provide important context to understand its worldwide invasion. The invasion of North America likely resulted from separate founding events on the East and West coasts (Commar et al. 2012). Comeault et al. (2020) showed that North American populations are genetically distinct from those from Africa. Invasive populations of *Z. indianus* have an approximately 30% reduction in genetic diversity relative to ancestral African populations (Comeault et al. 2020), though invasive populations of *Z. indianus* maintain levels of genetic diversity that are often higher than those of non-invasive congeners. Despite the loss of diversity, *Z. indianus* is extremely successful in temperate habitats (Rakes et al. 2023). Further studies demonstrated that genetically distinct populations from eastern and western Africa likely admixed prior to a single colonization of the Americas (Comeault et al. 2021). How the high degree of genetic diversity in invasive populations influences the potential for ongoing evolution in North America, which is in a critical early stage of invasion, remains understudied.

Here, we assembled and annotated a chromosome-level genome assembly for *Z. indianus* and used the newly improved genome to answer several questions with the whole genome sequences of over 200 North American flies collected from three locations over four years. First, do recolonizing North American *Z. indianus* populations demonstrate spatial or temporal population structure and if so, do specific regions of the genome have an outsized contribution to population structure? Second, is the invasion and recolonization history recapitulated in population genetic data? And third, do temperate populations show signatures of selection relative to native and tropical invasive populations?

## Materials and Methods

### Hi-C based genome scaffolding

An inbred line was generated from flies originally captured from Carter Mountain Orchard, VA (37.9913° N, 78.4721° W) in 2019. Wild caught flies were reared in the lab for approximately one year prior to initiating isofemale lines. The offspring of the isofemale lines were propagated through 10 rounds of full-sib mating. The resulting lines were then passaged for approximately one additional year in the lab and the most vigorous remaining line (“24.2”) was chosen for sequencing.

Third instar larvae from a single inbred line were snap frozen in liquid nitrogen and sent to Dovetail corporation (now Cantata Bio, Scotts Valley, CA) for chromatin extraction, Hi-C sequencing and genome scaffolding. Briefly, chromatin was fixed in place with formaldehyde in the nucleus and then extracted. Fixed chromatin was digested with DNase I, chromatin ends were repaired and ligated to a biotinylated bridge adapter followed by proximity ligation of adapter containing ends. After proximity ligation, crosslinks were reversed and the DNA purified. Purified DNA was treated to remove biotin that was not internal to ligated fragments. Sequencing libraries were generated using NEBNext Ultra enzymes and Illumina-compatible adapters. Biotin-containing fragments were isolated using streptavidin beads before PCR enrichment of each library. The library was sequenced on an Illumina HiSeqX platform to produce approximately 30x sequence coverage.

The input *de novo* assembly was the *Z. indianus* “RCR04” PacBio assembly (assembly # ASM1890459v1) from Kim et al. (2021). This assembly and Dovetail OmniC library reads were used as input data for *HiRise*, a software pipeline designed specifically for using proximity ligation data to scaffold genome assemblies (Putnam et al. 2016). Dovetail OmniC library sequences were aligned to the draft input assembly using *bwa* (Li and Durbin 2009). The separations of Dovetail OmniC read pairs mapped within draft scaffolds were analyzed by *HiRise* to produce a likelihood model for genomic distance between read pairs, and the model was used to identify and break putative misjoins, to score prospective joins, and make joins above a threshold. See Figure S1 for link density histogram of scaffolding data.

### Annotation

Repeat families found in the genome assemblies of *Z. indianus* were identified de novo and classified using the software package *RepeatModeler* v. 2.0.1 (Flynn et al. 2020). *RepeatModeler* depends on the programs *RECON* v. 1.08 (Bao and Eddy 2002) and *RepeatScout* v. 1.0.6 (Price et al. 2005) for the de novo identification of repeats within the genome. The custom repeat library obtained from *RepeatModeler* was used to discover, identify and mask the repeats in the assembly file using *RepeatMasker* v. 4.1.0 (Smit et al. 2015).

RNA sequencing was conducted on three replicates of 3^rd^ instar larva and three replicates of mixed stage pupa that were snap frozen in liquid nitrogen. RNA extraction and sequencing was performed by GeneWiz (South Plainfield, NJ). New larval and pupal RNAseq reads were combined with adult RNA sequencing from Comeault et al. (2020) for annotation. Coding sequences from *D. grimshawi, D. melanogaster*, *D. pseudoobscura*, *D. virilis*, *Z. africanus*, *Z. indianus*, *Z. tsacasi* and *Z. tuberculatus* (Kim et al. 2021) were used to train the initial *ab initio* model for *Z. indianus* using the *AUGUSTUS* software v. 2.5.5 (Keller et al. 2011). Six rounds of prediction optimization were done with the software package provided by *AUGUSTUS*. The same coding sequences were also used to train a separate *ab initio* model for *Z. indianus* using *SNAP* (version 2006-07-28) (Korf 2004). RNAseq reads were mapped onto the genome using the *STAR* aligner software (version 2.7) (Dobin et al. 2013) and intron hints generated with the *bam2hints* tools within *AUGUSTUS*. *MAKER* v. 3.01.03 (Cantarel et al. 2008), *SNAP* and *AUGUSTUS* (with intron-exon boundary hints provided from RNAseq) were then used to predict for genes in the repeat-masked reference genome. To help guide the prediction process, Swiss-Prot peptide sequences from the UniProt database were downloaded and used in conjunction with the protein sequences from *D. grimshawi, D. melanogaster, D. pseudoobscura, D. virilis, Z. africanus, Z. indianus, Z. tsacasi and Z. tuberculatus* to generate peptide evidence in the *MAKER* pipeline. Only genes that were predicted by both *SNAP* and *AUGUSTUS* were retained in the final gene sets. To help assess the quality of the gene prediction, AED scores were generated for each of the predicted genes as part of the *MAKER* pipeline. Genes were further characterized for their putative function by performing a BLAST search of the peptide sequences against the UniProt database. tRNA were predicted using the software *tRNAscan-SE* v. 2.05 (Chan and Lowe 2019). Transcriptome completeness was assessed with *BUSCO* v. 4.0.5 (Manni et al. 2021) using the eukaryota_odb10 list of 255 genes.

### Wild fly collections

Flies were collected by aspiration and netting from Carter Mountain Orchard, VA (in 2017-2020 and from Hanover Peach Orchard, VA in 2019-2020. Flies were sampled from Coral Gables, FL in June 2019 using traps baited with bananas, oranges, yeast, and red wine. See Table 1 for the number of individual flies sequenced from each location and timepoint. Flies were frozen in 70% ethanol at -20°C (2017-2018) or dry at -80 °C (2019-2020) prior to sequencing. Collections performed in July and August were called “early season.” In 2019, the earliest collections were not made until September (typically when *Z. indianus* abundance peaks, Rakes et al. 2023), and were assigned “mid-season.” Collections from October and November were called “late season.” For some analyses, the mid-season collection and early collections were combined, as they were the first collections available each year.

**Table 1.** Sample information for new samples collected for this study. See Table S1 for previously sequenced samples incorporated into this analysis.

| Location | Latitude, longitude | Year | Season | Sampling date (# flies) | Number sequenced | Number after filtering |
| --- | --- | --- | --- | --- | --- | --- |
| Coral Gables, FL, USA (FL) | 25.535, -80.493 | 2019 | - | 1 Jun | 25 | 24 |
| Carter Mountain, Virginia, USA (VA-CM) | 37.991, -78.472 | 2017 | early | 6 Jun (1)<br>22 Jun (7)<br>7 Jul (12) | 20 | 17 |
|  |  | 2017 | late | 9 Nov | 20 | 19 |
|  |  | 2018 | early | 5 Jul (2)<br>12 Jul (1)<br>19 Jul (1)<br>26 Jul (3)<br>2 Aug (1)<br>16 Aug (12) | 20 | 20 |
|  |  | 2018 | late | 1 Nov | 20 | 19 |
|  |  | 2019 | mid | 6 Sep | 20 | 20 |
|  |  | 2019 | late | 26 Oct | 20 | 14 |
|  |  | 2020 | early | 17 Jul | 20 | 18 |
| Hanover Peach Orchard, Virginia, USA (VA-HPO) | 37.572, -77.266 | 2019 | mid | 3 Sep | 20 | 19 |
|  |  | 2020 | early | 16 Jul (5)<br>3 Aug (25) | 30 | 29 |

### Individual whole genome sequencing

The sex of each wild-caught fly was recorded, then DNA was extracted from individual flies using the DNAdvance kit (Beckman Coulter, Indianapolis, IN) in 96 well plates, including an additional RNase treatment step. DNA concentration was measuring using the QuantIT kit (Invitrogen, Waltham, MA) and purified DNA was diluted to 1 ng/µL. Libraries were prepared from 1 ng of genomic DNA using a reduced-volume dual-barcoding Nextera (Illumina, San Diego, CA) protocol as previously described (Erickson et al. 2020). The libraries were quantified using the QuantIT kit and equimolar ratios of each individual DNA were combined for sequencing. The pooled library was size-selected for 500 bp fragments using a BluePippin gel cassette (Sage Sciences, Beverly, MA). The pooled libraries were sequenced in one Illumina NovaSeq 6000 lane using paired-end, 150 bp reads by Novogene (Sacramento, CA).

Existing raw reads from *Z. indianus* collections from North America, South America, and Africa (Table S1; Comeault et al. 2020; Comeault et al. 2021) were downloaded from the SRA from BioProject number PRJNA604690. These samples were combined with the new sequence data and processed together with the same mapping and SNP-calling pipeline. Overlapping paired-end reads were merged with *BBMerge* v. 38.92 (Bushnell et al. 2017). Reads were mapped to the genome assembly described above using *bwa mem* v. 0.7.17 (Li and Durbin 2009). Bam files for merged and unmerged reads were combined, sorted and de-duplicated with *Picard* v. 2.26.2 (https://github.com/broadinstitute/picard).

We next used *Haplotype Caller* from *GATK* v. 4.2.0.0 (McKenna et al. 2010) to generate a gVCF for each individual. We built a GenomicsDBI database for each scaffold, then used this database to genotype each gVCF. We used *GATK’s* hard filtering options to filter the raw SNPs based on previously published parameters (*--filter-expression “QD < 2.0 || FS > 60.0 || SOR > 3.0 || MQ < 40.0 || MQRankSum < -12.5 || ReadPosRankSum < -8.0”*) (Comeault et al. 2020). We then removed SNPs within 20 bp of an indel from the output and removed all SNPs in regions identified by *RepeatMasker*. We analyzed several measures of individual and SNP quality using *VCFtools* v. 0.1.17 (Danecek et al. 2011). We removed 16 individuals with mean coverage < 7X or over 10% missing genotypes. Next, we filtered SNPs with mean depths <10 or > 50 across all samples. We removed individual genotypes supported by 6 or fewer reads or with more than 100 reads to produce a final VCF with 5,185,389 SNPs and 2,099,147 non-singleton SNPs. See Table 1 and S1 for the final number of individuals included in the analysis from each population. See Figure S2 for the average SNP depth per sampling time and location.

### Sex chromosome and Muller element identification

*samtools* v. 1.12 (Li et al. 2009) was used to measure coverage and depth of mapped reads from individual sequencing. This analysis revealed that the five main scaffolds (all over 25 Mb in length) had a mean depth of ∼16X coverage in both males and females in our dataset, except for scaffold 3, which had ∼16X coverage in females but ∼8X coverage in males, suggesting it is the X chromosome (Figure S3). Some of the previously sequenced samples had no sex recorded, so we used the ratio of X chromosome reads (scaffold 3) to autosome (scaffolds 1, 2, 4 and 5) reads to assign sexes to those individuals. Individuals with a ratio greater than 0.8 were assigned female, and ratios less than 0.8 were assigned male (Figure S4). For two known-sex individuals, the sex recorded prior to sequencing did not match the sex based on coverage; for those two samples we used the coverage-based sex assignment for analyses. We used *D-GENIES* (Cabanettes and Klopp 2018) to create dot-plots comparing the *Z. indianus* and *D. melanogaster* genome (BDGP6.46, downloaded from ensemble.org) to confirm the sex chromosome identification and assign Muller elements to *Z. indianus* autosomes (Figure S5, Table S2). Five additional scaffolds had lengths over 1 Mb. Scaffold 8 is the dot chromosome (Muller element F) based on sequence comparison to *D. melanogaster* (Figure S5) and had similar coverage to the autosomes (Figure S3). Scaffolds 6,7,9, and 10 had reduced coverage (Figure S3) and contain mostly repetitive elements. Downstream SNP calling and population genetic analysis included the five large scaffolds (named chromosomes 1-5 from largest to smallest) and excluded all smaller scaffolds.

### Testing for structural variants

We used *smoove* v. 0.2.6 (Pedersen et al. 2020) to identify and genotype insertions, deletions, and rearrangements in the paired-end sequencing data from all individuals as described in the documentation. As an alternative approach to search for large structural variants, we used linkage disequilibrium (LD) of randomly sampled SNPs from each chromosome to visually inspect for linkage due to potential inversions. We generated a list of SNPs segregating in each focal population with no missing genotypes and randomly sampled 4,000 SNPs from each chromosome. We used the *snpgdsLDMat* function in *SNPRelate* to calculate LD between all pairs of SNPs. LD heatmaps were created with the *ggLD* package (https://github.com/mmkim1210/ggLD).

To define approximate inversion regions for SNP filtering, we looked for regions with evidence of high long-distance linkage disequilibrium within North American samples. We note that our purpose here was to roughly define inversion regions to exclude from population structure analyses, not to precisely define inversion breakpoints, which will likely require further long-read sequencing. For each SNP segregating in North America with complete genotyping information, we randomly chose a second SNP that was 100, 200, 300, 400, or 500 kb away from the focal SNP (+/- 5%). We calculated LD between the pair of SNPs. We then divided the genome in 100 kb nonoverlapping windows and for each window determined if at least one SNP in that window showed evidence of high long-distance LD (R^2^ > 0.75) at any of the sampled distances. Because some windows contain spurious examples of long-distance LD, we defined inversion regions by looking for consecutive strings of at least 10 windows (1 Mb total) containing high LD SNPs. This approach identified large potential inversion regions on chromosomes 1, 2, 4, and 5 that corresponded to regions visually identified in LD heatmaps. We masked SNPs in these regions (82.6 Mb total) for some downstream analyses.

### Population structure

We conducted principal components analysis (PCA) using the R package *SNPRelate* v. 1.38.0 (Zheng et al. 2012) in R v. 4.1.1 (R Core Team) using a vcf that excluded singleton SNPs and potential inversion regions. We LD pruned SNPs with minor allele counts of at least 3 (Linck and Battey 2019) using *SNPgdsLDpruning* with an LD threshold of 0.2 and then calculated principal components with *snpgdsPCA* using all four autosomes. For subsequent analyses, we repeated the LD pruning within subsets of the data (North America only, or Carter Mountain, VA only). We also calculated principal components using individual chromosomes; for the X chromosome, only females were used in the analysis. The single fly collected in Kenya 2018 (Comeault et al. 2021) was an extreme outlier in the preliminary PCA and was excluded. We conducted the initial analysis using all SNPs that met the filtering criteria and then conducted a second analysis using SNPs that were masked for potential inversion regions as described above. We conducted discriminant analysis of principal components using the *dapc* function in package *adegenet* 2.1.10 (Jombart 2008; Jombart and Ahmed 2011). We used the *optim.a.score* function to determine the number of PCs to include in the DAPC analysis.

We used *Plink* v. *1.9* (Purcell et al. 2007; Chang et al. 2015) to LD prune VCF files with parameters (*--indep-pairwise 1000 50 0.2*) and used *ADMIXTURE* v. 1.3.0 (Alexander and Lange 2011) to evaluate population structure. Using whole genome data not masked for inversions, we calculated admixture for each chromosome separately. For the X chromosome, only females were used. We tested up to *k*=10 genetic clusters and used cross-validation analysis to choose the optimal *k* for each chromosome separately. We also ran ADMIXTURE using all non-inversion autosomal SNPs together.

We calculated F_ST_ between Virginia samples and those from Florida or Africa using the *snpgdsFST* function in *SNPRelate* for all SNPs with a minor allele count > 3. For the X chromosome, only females were used in F_ST_ calculations. We used the same function to calculate genome-wide, pairwise F_ST_ between all Virginia collections using autosomal SNPs with putative inversions masked. To quantify relatedness between individuals, we used the function *snpgdsibdKING* in *SNPRelate* v. 1.38.0 (Zheng et al. 2012) to determine the kinship coefficients and probability of zero identity by descent for pairs of individuals using autosomal SNPs with putative inversions masked. We used thresholds established in Thornton et al. (2012) to classify relatedness between individuals. We used the *–het* function in *vcftools 0.1.17* to calculate the inbreeding coefficient for each individual using non-inversion SNPs.

We generated phylogenetic trees using *Treemix* (Pickrell and Pritchard 2012). We generated a vcf masked for inversions that contained only Carter Mountain and Florida samples. We LD-pruned this vcf in PLINK 1.9 with *--indep-pairwise 100 10 0.2* and prepared *Treemix* input files with custom scripts. We then ran 100 bootstrap replicates of *Treemix* with zero migration edges, Florida as the outgroup, and arguments *–bootstrap -k 500.* We used *PHYLIP* v. 3696 (Felsenstein 2005) to generate a consensus tree and support for each node. We plotted a representative output in R using scripts from https://github.com/andrewparkermorgan/popcorn/.

### Estimation of historic population sizes

We used *smc*++ v. 1.15.4 (Terhorst et al. 2017) to estimate historic population sizes for several subpopulations of individuals using chromosome 4 genotypes since this chromosome lacked evidence of large structural rearrangements. We used individuals from each African location and used the earliest sampling available for each year and Virginia orchard. We used *vcf2smc* to prepare the input files for each autosome separately. We assigned each individual as the “distinguished individual” and ran the analysis using all possible combinations of distinguished individual as described in (Bemmels et al. 2021). We used cross validation to estimate final model parameters with the option (*-cv –folds = number of individuals)*. We assumed a generation time of 0.08 years (∼12 generations per year) based on Nava *et al*. (2007), which assumes year-round reproduction in tropical regions. We note that for Virginia populations experiencing temperate conditions in recent years, 12 generations per year is likely an overestimate due to the shortened breeding season.

### Selection scan

We used *WhatsHap* v. 1.7 (Patterson et al. 2015) to perform read-based phasing of the full vcf including singletons. To polarize the vcf for the genome wide selection scan relative to the invasion, we reassigned the reference allele of the phased vcf as the allele that was most common across all African individuals sequenced in previous studies. We calculated allele frequencies using all African samples in *SNPRelate*, then used *vcf-info-annotator* (https://vatools.readthedocs.io/en/latest/index.html) to assign the “ancestral” allele in the INFO column. Lastly we used *bcftools* v. 1.13 (Danecek et al. 2021) to make simplified vcfs containing only the GT and AA fields for each chromosome separately.

We used the R package *rehh* v. 3.2.2 (Gautier and Vitalis 2012) to conduct the selection scan using integrated haplotype homozygosity score (IHS). The test measures the decay of haplotype homozygosity to look for long, shared haplotypes that are signatures of selective sweeps when a single haplotype rises to high frequency without being eliminated by recombination (Sabeti et al. 2002; Voight et al. 2006). We used all flies from Virginia and conducted the scan using phased, polarized vcfs for each individual chromosome. We used the *haplo2hh*, *scan,* and *ihh2ihs* functions to implement the scan. For the X chromosome, we only used a single haplotype for each male in the dataset to avoid double counting haploid genotypes.

We also used *BayPass v. 2.41* (Gautier 2015) to calculate signals of selection comparing flies from Virginia to those from Florida and Africa while accounting for population structure via the XtX statistic. For this analysis, we used only females so that a single analysis could be conducted across all chromosomes. We created separate vcf files for each population and prepared *BayPass* input files using *vcftools –counts2*. We pruned these files for linkage disequilibrium in *Plink* 1.9 using *--indep-pairwise 50 5 0.2.* We then ran *BayPass* according to the user manual and modified code from (Whiting et al. 2021) to generate a core model and covariance matrix and final XtX values. We used 10,000 simulated SNPs to define XtX outliers.

To look for shared signals across the F_ST_, XtX and IHS selection/differentiation analyses, we used a window-based approach. We divided the genome into 1kb windows with a 500 bp step size and identified all windows that contained at least one SNP that fell in the top 99% of SNPs for the F_ST_, XtX and IHS tests.

### Genetic diversity statistics

Because we obtained variable sequencing coverage within and across populations (Figure S2) we used software designed for low coverage and missing data to analyze population genetic statistics in genomic windows. We used *pixy* v. 1.2.5 (Korunes and Samuk) to calculate Pi, F_ST_ and D_XY_ in 5 kb windows. Samples were grouped by collection location; only females were used for analysis of the X chromosome. We used *ANGSD* v. 0.941 (Korneliussen et al. 2014) to calculate Tajima’s D. We first calculated genotype likelihoods from the bam files using arguments *-doSaf* and *-GL*. We then calculated Tajima’s D and theta using the folded site frequency spectrum across 5 kb windows with 5 kb steps as described in *ANGSD* documentation using only female samples.

### Data management and plotting

We used the R packages *foreach* (Microsoft and Weston 2017) and *data.table* (Dowle and Srinivasan 2019) for data management and manipulation and used *ggplot2* (Wickham 2016) for all plotting. The *ggpubfigs* (Steenwyk and Rokas 2021) and *viridis* (Garnier 2018) packages were used for color palettes.

## Results and Discussion

### Genome assembly and annotation

High quality genome assemblies and annotations are a critical component of tracking and controlling invasive species and understanding their potential for evolution in invaded ranges (Matheson and McGaughran 2022). We conducted Hi-C based scaffolding of a previously sequenced *Z. indianus* genome (Kim et al. 2021) to achieve a chromosome-level assembly. There were 1,014 scaffolds with an N50 of 26.6 Mb, an improvement from an N50 of 4.1-6.8 Mb in previous assemblies (Kim et al. 2021). The five main chromosomes (Figure S1, named in order of size from largest to smallest) varied in length from 25.7 to 32.3 Mb (total length of five main scaffolds = 146,062,119 bp), in agreement with *Z. indianus* karyotyping (Gupta and Kumar 1987; Campos et al. 2007). Chromosome 3 was identified as the X chromosome using sequencing coverage of known-sex individuals (Figure S3, S4). See Table S2 for assignment of *Z. indianus* chromosomes to Muller elements based on alignment to the *D. melanogaster* genome (Figure S5).

The annotation using RNAseq from larvae, pupae, and adults predicted 13,162 transcripts and 13,075 proteins, with 93% of 255 benchmarking universal single copy orthologs (BUSCO) genes (Simão et al. 2015) identified as complete and an additional 1.2% of BUSCO genes identified as fragmented. This transcriptome-based completeness estimate is lower than the genome-based estimate of 99% complete (Kim et al. 2021) but is in line with other arthropod genomes (Feron and Waterhouse 2022). Within the five main chromosomes, 24.6% of sequences were repetitive; within the entire assembly including all smaller scaffolds, 41% were repetitive. The five main chromosomes contain 11,327 predicted genes (87% of all predicted), including 99.5% of all complete BUSCO genes. This improved genome resource will be valuable for future evolutionary studies *of Z. indianus*, which is becoming an increasingly problematic pest in some regions of the world (Allori Stazzonelli et al. 2023).

### Limited spatial or temporal population structure in North American Z. indianus

To study spatial and temporal patterns of genetic variation in the seasonally repeated invasion of *Z. indianus*, we resequenced ∼220 individuals collected from two orchards in Virginia (Carter Mountain Orchard and Hanover Peach Orchard) from 2017-2020, as well as one population collected from Coral Gables, Florida in 2019. Because temperate locations such as Virginia are thought to be recolonized by southern populations of *Z. indianus* each year (Pfeiffer et al. 2019; Rakes et al. 2023), we sampled both early in the season (∼July-August) and late in the season (∼October-November) in each year to capture the founding event, population expansion, and potential adaptation to the temperate environment.

We were first interested in studying geographic and temporal variation in population structure in North American populations of *Z. indianus*. For this analysis, we incorporated previous sequencing data from the Western Hemisphere and Africa (Table S1 and Comeault et al. 2020). While previous studies have shown limited structure within North America (Comeault et al. 2020; Comeault et al. 2021), we wanted to test for structure using deeper sampling within introduced locations and with greater temporal resolution across the *Z. indianus* growing season (Rakes et al. 2023). Our initial analyses of population structure using principal components analysis (PCA) of the whole genome (Figures S6a) and individual chromosomes (Figure S7a) and *ADMIXTURE* (Alexander and Lange 2011) analysis of individual chromosomes (Figure S8) confirmed previous results that Western Hemisphere samples are genetically distinct from African samples. To focus on potential structure within invasive North American samples, we excluded the African samples and recalculated principal components. North American samples fell into three large groups when considering PC1 and PC2 (diagonal bands in Figure S6B), but these clusters generally did not correspond to sampling locations. In single-chromosome PCA, North American samples formed clusters that did not correspond to geographic locations for chromosomes 1, 2, 3, and 5; such patterns are potentially indicative of inversions that influence genotypes of many SNPs simultaneously (Li and Ralph 2019). This pattern was upheld in the *ADMIXTURE* analysis; notably, for chromosomes 1, 2, and 5, many individuals showed ∼50% ancestry assignment to different clusters, which could reflect genotypes for large structural rearrangements. These findings led to further investigations of potential structural polymorphisms (described below) and the masking of SNPs potentially located in large chromosomal rearrangements to examine population structure.

After removing regions of the genome that were potentially part of inversions (see below), PC1 still separated Western Hemisphere and African samples (Figure 1a), and North American samples remained tightly grouped for PCs 1-4, indicating most genetic variation is found within African samples. Analysis of only North American samples revealed little genome-wide separation of populations collected from different locations (Figure 1b), though some Carter Mountain, Virginia individuals were outliers from the main cluster of points. Many invasive species evolve complex population structures in the invaded range due to a combination of bottlenecks, founder effects and rapid local adaptation (Koch et al. 2020; Atsawawaranunt et al. 2023; García-Escudero et al. 2023). On the other hand, some invasive species have more homogenous populations across widespread invaded ranges in eastern North America (Friedline et al. 2019; Barrett et al. 2023). A high rate of migration between orchards (occurring naturally or due to human-mediated transport) or large founding population sizes could result in a lack of geographic differentiation between populations.

**Figure 1:**
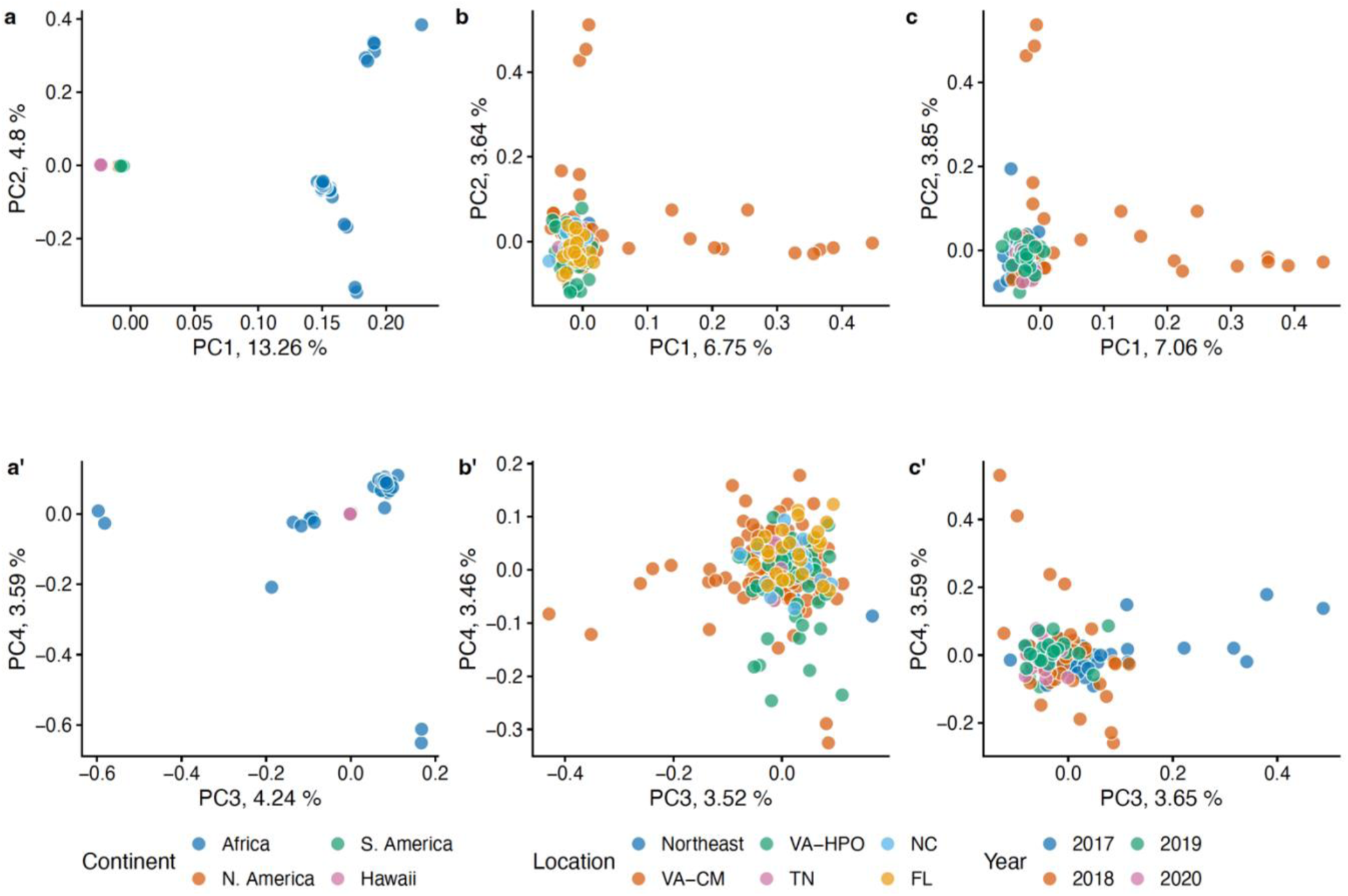
Principal component analysis of Z. indianus populations using autosomal SNPs outside of putative inversions reveals a lack of spatial or temporal population structure in invasive populations. Top row shows PC1 and PC2; bottom row shows PC3 and PC4. (a-a’) All individuals (n=266), color coded by continent/locale of collection. South America, North America, and Hawaiian samples are all tightly clustered such that individual points are not visible. (b-b’). All North American individuals (n=224), color coded by collection site; HPO and CM are two orchards in Virginia; Northeast refers to samples from NY, NJ, and PA. (c-c’) All individuals from Carter Mountain, Virginia (n=127), color coded by year of collection. For each analysis, only the individuals shown in the plot were included in the PC calculation.

We next hypothesized that founder effects during each recolonization event might lead to unique genetic compositions of temperate populations sampled in different years (Uller and Leimu 2011). We calculated principal components using only samples collected from Carter Mountain, VA in 2017-2020. Surprisingly, in these samples, we saw no evidence of population structure between years (Figure 1c), apart from some samples from 2018 that were divergent from the rest; we investigate potential causes of this pattern below. These data suggest that the founding fly populations in Virginia are relatively homogeneous each year at a genome-wide scale, but some years may contain genetically divergent individuals. This result is consistent with the lack of spatial population structure and likewise could indicate large founding populations or ongoing migration. Alternatively, the Virginia populations could be permanently established with little genetic differentiation year-to-year, though this possibility is not supported by field data from temperate locations (Rakes et al. 2023; Kohlmeier and Kohlmeier 2025).

*ADMIXTURE* revealed two ancestral clusters for global *Z. indianus* populations, separating Africa from North America (Figure 2). Adding increasing numbers of ancestral groups revealed genetic differentiation within North American samples, but this differentiation did not correspond to location or year of collection. Notably, with k=3, some individuals from Carter Mountain in 2018 separated from the other North American samples; these samples correspond to those that were outliers in the PCA. Collectively, our results are consistent with previous findings that African populations are distinct from those in the Western Hemisphere, and we find little evidence of genome-wide population structure over space or time within North American samples.

**Figure 2:**
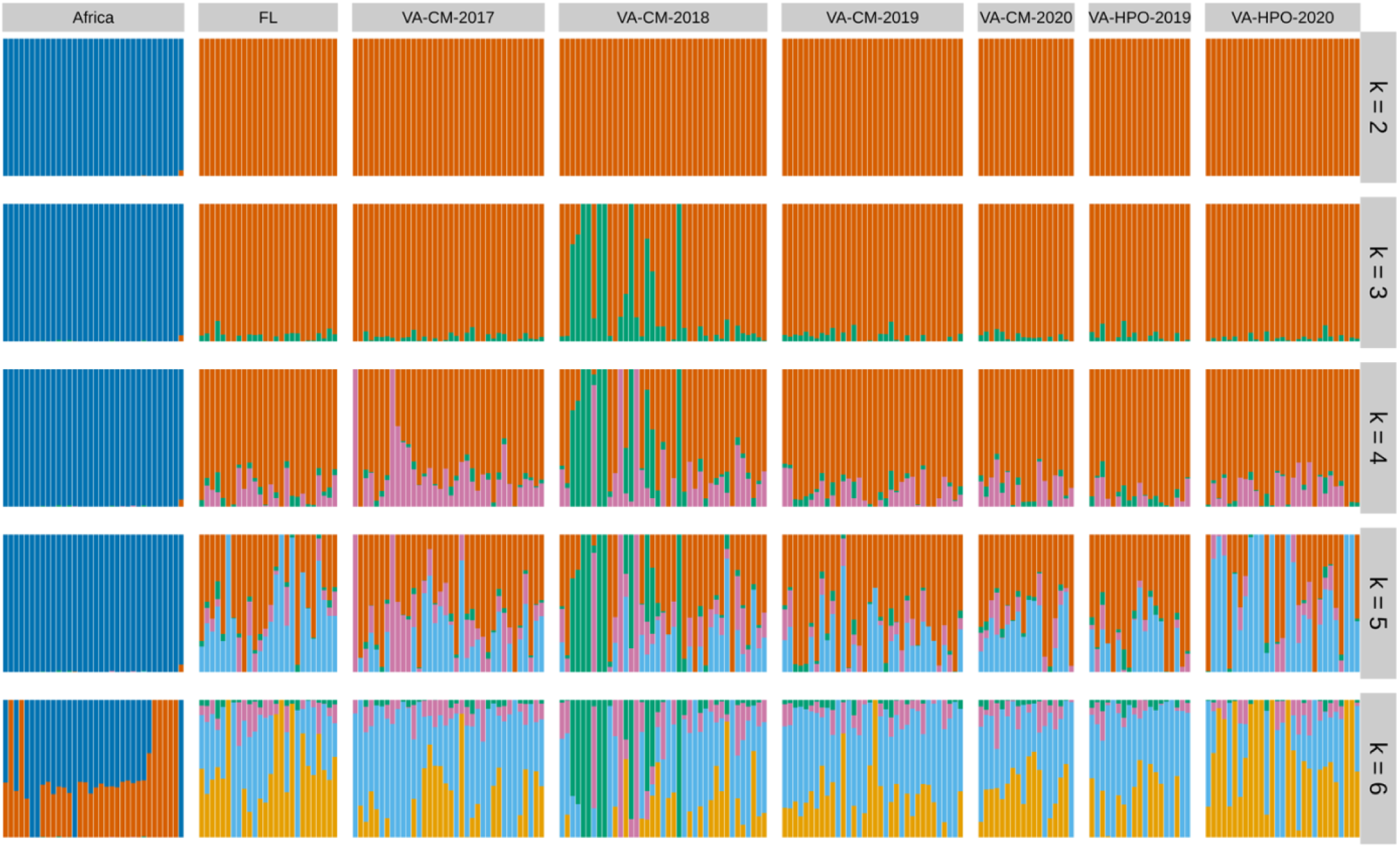
Admixture analysis of Z. indianus from different locations. Each column is an individual, and colors represent assignment to distinct genetic clusters based on all autosomal SNPs outside of inversion. Each row represents a different number of ancestral genetic clusters (k = 2 to 6). FL=Coral Gables, Florida, VA-HPO = Hanover Peach Orchard, Virginia, VA-CM = Carter Mountain, Virginia. African sequences combine five geographic locations from Comeault et al. (2020 & 2021). Cross-validation analysis supported k=2 (top row) as the most likely number of ancestral groups.

### Structural polymorphism

The clustering of samples in the single-chromosome PCA (Figure S7), combined with many individuals showing ∼50% assignment to genetic clusters within individual chromosomes (Figure S8), suggested that large structural variants may be segregating in *Z. indianus* (Li and Ralph 2019; Nowling et al. 2020). To look for evidence of structural variants via depressed recombination rates, we examined linkage disequilibrium (LD) from 4,000 randomly sampled SNPs on each chromosome. In North American samples, we discovered large blocks of LD spanning substantial portions of chromosomes 1, 2, 3, and 5 (Figure S9), potentially indicative of inversions (Fang et al. 2012; da Silva et al. 2019). However, there was no evidence of long-distance LD in these regions in the African samples (Figure S9). These results suggest inversions on chromosomes 1, 2, 3, and 5 are segregating in North America but are relatively rare in Africa. To approximately define the boundaries of these inversions, we calculated LD for random pairs of SNPs that were 100-500 kb apart on each chromosome. This analysis similarly identified large regions containing elevated long-distance LD on chromosomes 1, 2, 4, and 5 (Figure S10A). We compared the results of the LD analysis to potential inversions called using paired-end data by *smoove* but found little correspondence (Figure S10B, compare gray boxes to yellow boxes). As such, we use the LD-based data to mask potential inversion regions for population structure analyses described above.

Given the relative chromosome sizes in the genome assembly, the linkage blocks on chromosomes 2 and 5 likely correspond to the previously described inversions *In(V)B* and *In(II)*A, respectively (Ananina et al. 2007). Since chromosome 1 is the longest chromosome in our assembly, the linkage likely correspond to the complex *In(IV)EF* polymorphism, made up of two overlapping inversions (Ananina et al. 2007); different genotypic combinations of two inversions could explain the six distinct clusters seen in the chromosome 1 PCA (Figure S7b). The X chromosome has three described inversions in *Z. indianus* (Ananina et al. 2007), which may explain to the complex pattern of linkage observed in North American samples on this chromosome (Figure S9, S10) and the clustering of North American samples in the chromosome 3 PCA (Figure S6B,C). Major chromosomal polymorphisms are known to be important for local adaptation and phenotypic divergence in a wide variety of species (Joron et al. 2011; Küpper et al. 2016; Lee et al. 2016; Huang et al. 2020; Nunez et al. 2024), including inversions that facilitate invasive phenotypes (Galludo et al. 2018; Tepolt and Palumbi 2020; Tepolt et al. 2022; Ma et al. 2024). These inversions may be new to North America, or they may have been present at low frequency in the bottlenecked population that founded *Z. indianus* populations in the Western Hemisphere but then experienced subsequent selection in the invaded range. Alternatively, they have arisen in a currently unsequenced population and then been introduced to the Western Hemisphere. Further characterization of these inversions via additional sequencing and phenotypic characterization to determine whether they influence *Z. indianus*’ survival or fitness in the invaded range will be a rich area for future studies.

### Recolonization, bottlenecks and seasonal dynamics in Z. indianus

Invasive species typically experience a genetic bottleneck due to small founding population sizes (Barrett 2015; Estoup et al. 2016). We hypothesized that North American populations would show reduced effective population size (N_e_) relative to African populations, and that Virginia populations would show a further, more recent reduction in N_e_ relative to Florida populations as the result of a secondary population bottleneck upon temperate recolonization. We estimated historic population sizes using *smc++* (Terhorst et al. 2017) using data only from chromosome 4, which lacks large inversions (Figure 3a). Our prediction was correct with respect to Africa vs North America: African populations show historical fluctuations but population sizes typically in the range of ∼ 10^4^-10^7^ individuals (10^6^-10^7^ for the past ∼1,000 years). However, introduced populations in North America demonstrated a dramatic reduction in population size in the past ∼100 years, perhaps reflecting a bottleneck following colonization of Brazil in the late 1990s (Yassin et al. 2008) and consistent with the previously described loss of genetic diversity in invasive populations (Comeault et al. 2020). This contraction is followed by a rebound as introduced populations expanded over the past several decades. Overall, the ancestral population sizes for Virginia and Florida were quite similar with overlapping ranges of the estimates, and our prediction of reduced recent population sizes in Virginia relative to Florida was not well-supported. Given our limited sample sizes and potential differences in the number of generations per year in temperate and subtropical environments, detecting fine-scale differences in very recent population fluctuations may be beyond the detection ability of the software; *smc++* becomes less accurate at timescales less than ∼333 generations (∼26 years for year-round populations of *Z. indianus*) (Patton et al. 2019). Virginia populations could be colonized by a large number of individuals, or they may represent admixed populations reflecting individuals from multiple sources, producing larger effective population sizes than would otherwise be expected if recolonization occurs from a single source population undergoing a bottleneck. Admixture and gene flow are important factors fueling genetic diversity and invasiveness in introduced species (McGaughran et al. 2024) and could potentially contribute to *Z. indianus’* local success following each recolonization event.

**Figure 3:**
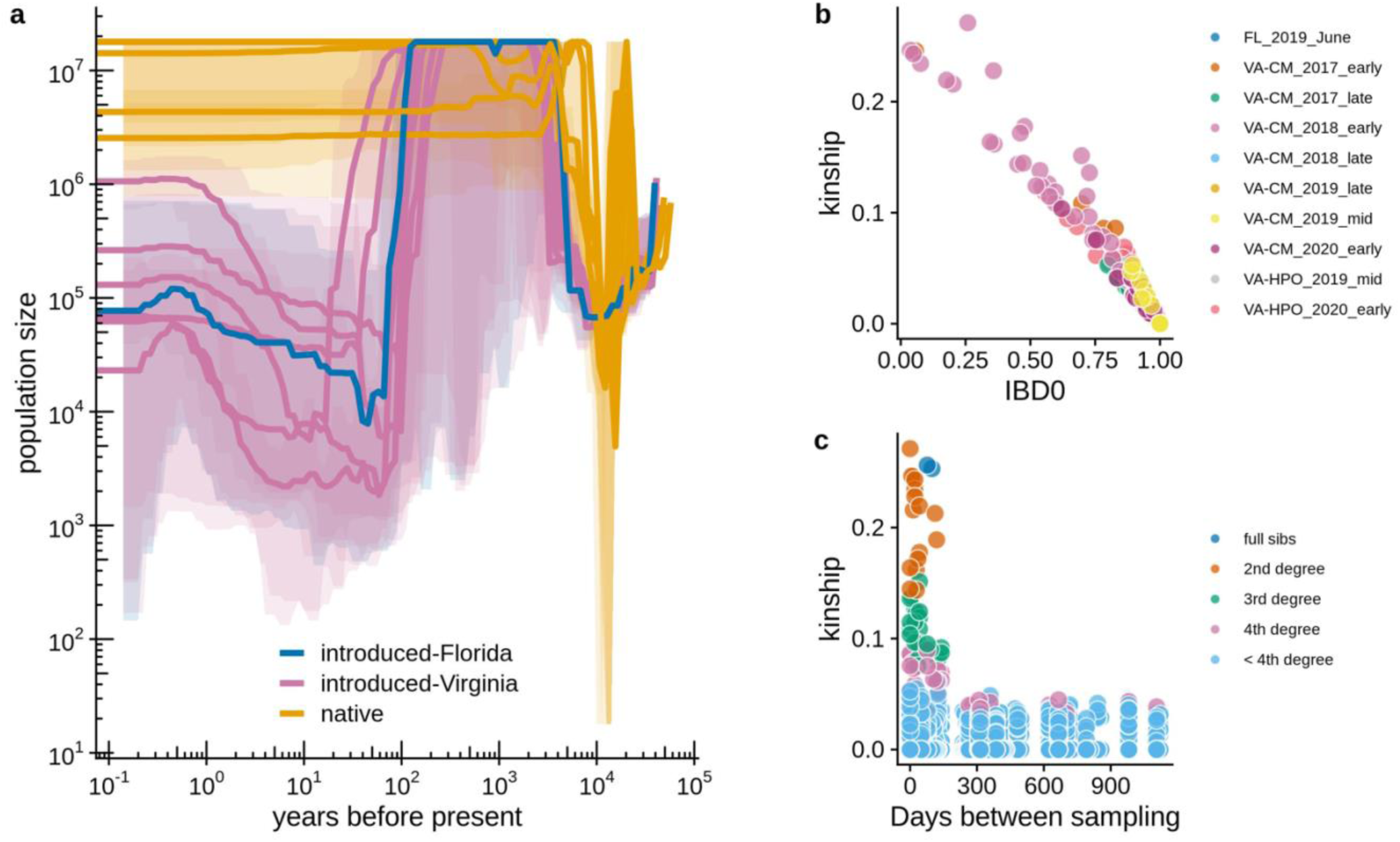
Demographic effects of bottlenecks in Z. indianus populations. a) Population history reconstruction with smc++ using genotypes from chromosome 4, which lacks inversions. Introduced-Virginia flies were collected in the early-mid season (June-September) from two Virginia orchards in 2017-2020 (n=6 populations grouped by orchard and year). Native populations are five distinct African populations (Kenya, Zambia, Senegal-Forest, Senegal-Desert, and Sao Tome [Comeault et al. 2020]). Shaded ribbons show minimum and maximum population sizes estimated in cross-validation analysis. b) Kinship and probability of zero identity by descent for pairs of individual flies from the same collection location and season within North America calculated with non-inversion autosomal SNPs. c) Kinship coefficients for pairs of individual flies collected at Carter Mountain Orchard, Virginia, as a function of the number of days between sampling. Relatedness was assigned according to thresholds from Thornton et al. (2012).

We additionally tested for bottlenecks by looking for the presence of relatives within our samples, which might be a product of small founding populations. Using two measures of genetic similarity, we discovered many pairs of related flies in our dataset (Figure 3B). Most dramatically, many flies collected in 2018 appeared to be close relatives (Figure S11), consistent with the separation of some 2018 samples in the PC and *ADMIXTURE* analyses (Figures 1c and 2). In collections from late July and early August 2018, 26 pairs of close relatives involving 13 individual flies were collected. Of those, 21 pairs of relatives were collected on different days, suggesting the relatedness was not solely a sampling artifact due to collecting relatives in the same microhabitat of the orchard. The effect of this apparent bottleneck was sometimes retained throughout the growing season, as a pair of full sibs was sampled 77 days apart in 2018, two pairs of second-degree relatives were sampled over 110 days apart in 2018, and two pairs of third-degree relatives were sampled 140 days apart in 2017 (Figure 3c). Given that *Z. indianus* are collected in small numbers early in the season (Rakes et al. 2023) and 2017 and 2018 had particularly early captures (Table 1), we thought a small founding population size followed by inbreeding could produce individuals sampled distantly in time that still show close genetic similarity. However, we did not find any significant difference in inbreeding coefficients between flies sampled at different time points at Carter Mountain (ANOVA, F(6,120) = 1.79, P = 0.106). Instead, flies may live for a relatively long time or have slower generations in the wild, allowing us to capture close relatives separated by longer time periods. However, we note that the same pattern was not seen in every year of our collections, suggesting that colonization dynamics might differ dramatically from year to year, which is expected if recolonization occurs due to chance events each year.

A founder effect could generate temporal population structure by creating populations that were more similar within a year than between years, creating a positive relationship between F_ST_ and the elapsed time between collections (Bergland et al. 2014). We tested this prediction with samples collected from Carter Mountain, Virginia over four years and found a weak, positive correlation between F_ST_ and the time between sample collections for comparisons that excluded the unusual 2018 collection containing many relatives (linear model, df=19, R^2^ = 0.423, P=0.009, Figure S12). This finding is consistent with trends observed in *D. melanogaster,* which experiences a strong overwintering bottleneck and shows temporal patterns of differentiation (Bergland et al. 2014; Nunez et al. 2024). Since this finding could be caused by an overwintering bottleneck or a recolonization bottleneck, we further investigated population relationships over time.

We used discriminant analysis of principal components (DAPC) and phylogenetic trees to look at relationships between flies captured at Carter Mountain, Virginia, over time relative to the population from Florida. In North America, *Z. indianus* was first found in Florida and was later found northwards (Linde et al. 2006; Pfeiffer et al. 2019). If flies are permanently established in Virginia, we predict that the Virginia populations would become progressively more different from Florida over time in a “stepping-stone” like pattern as each population evolves independently. If flies recolonize each year with small founding population sizes, we expect populations to be more similar to Florida in some years than others due to chance. The predictions of recolonization were supported by the DAPC analysis, which showed that the 2019 sample shared the most overlap with Florida, while 2018 was the most different (Figure 4a), and there was no clear pattern of temporal differentiation. We also used *Treemix* (Pickrell and Pritchard 2012) and *PHYLIP* (Felsenstein 2005) to build a consensus phylogenetic tree from 100 independent bootstraps of the same dataset. The topology of the consensus tree (Figure 4b) was not consistent with stepwise divergence of the populations over time; the two earlier timepoints formed a clade and the two later timepoints formed a clade, though the node for the 2019-2020 divergence was not well supported. The drift parameter was greater in 2017 and 2018 than it was 2019 and 2020, suggesting that populations collected later in time were more similar to Florida than those collected earlier in time. This result is consistent with a recolonization model in which the founding population is randomly more representative of the source population in some years than others, with smaller and earlier founding events in 2017 and 2018 potentially leading to greater genetic drift. Additional demographic simulations will likely be required to more definitively determine the colonization dynamics of *Z. indianus*.

**Figure 4:**
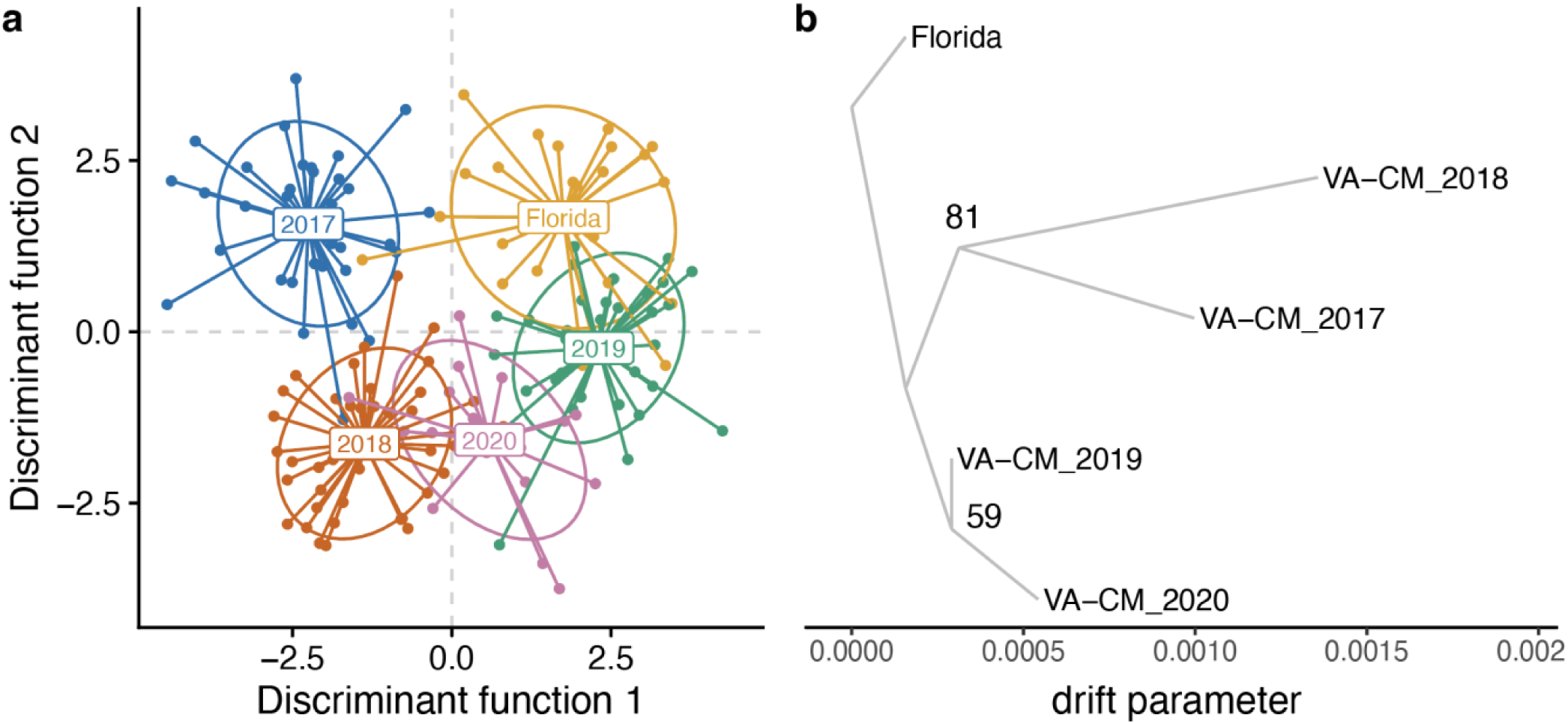
DAPC and Treemix analyses suggest Z. indianus recolonizes Virginia each year. All analyses were conducted with autosomal SNPs outside of inversions. a) Discriminant analysis of principal components (DAPC) of samples from Florida and four years of sampling from Carter Mountain, Virginia. b) Results of a single Treemix run representative of the consensus tree topology from PHYLIP; bootstrap support values from 100 trees are given at nodes.

### Signals of recent selection in Z. indianus near potential pesticide resistance genes

We tested for signals of recent natural selection in temperate *Z. indianus* populations from Virginia. We compared Virginia populations to those from Africa to test for broader invasive-native population differentiation and compared Virginia to Florida to look for potentially more recent temperate adaptation. We measured three statistics: SNP-level F_ST_ comparing all flies collected Virginia to those from Florida or Africa; the XtX statistic from *BayPass* (Gautier 2015) to test for selection in the presence of population structure, and integrated haplotype homozygosity score (IHS) to look for long, shared haplotypes that are signatures of recent selective sweeps when a single favorable haplotype rises to high frequency (Sabeti et al. 2007). We note that our approach would not detect sweeps involving multiple alleles from standing variation (soft sweeps; Messer and Petrov 2013; Garud et al. 2015), which could be an important potential component of *Z. indianus* evolution given the high levels of genetic diversity found in invasive populations (Avalos et al. 2017; Comeault et al. 2020). To compare results across tests, we looked for 1 kb windows containing SNPs that fell in the top 1% of all three tests. As expected given the high degree of population structure, SNPs with high F_ST_ between Africa and Virginia were widespread across all chromosomes, and F_ST_ was particularly high on the X chromosome (Figure 5a). This observation is in line with the findings of Comeault et al. (2021), who showed that many X-linked scaffolds showed signs of selection in invasive populations, and we suggest this pattern is likely related to the smaller effective population size for X chromosomes (Ellegren 2009) and the presence of several inversions on this chromosome (this study; Ananina et al. 2007). After accounting for population structure with *BayPass*, the signal remained high throughout most of the X chromosome, but several peaks of potential selection were resolved on the autosomes, including prominent peaks on chromosomes 2 and 5 (Figure 5c). These peaks corresponded to windows with long, shared haplotypes in Virginia populations (high IHS scores, Figure 5e). The complex population structure and broadly elevated signals of divergence and selection on the X chromosome made it difficult to investigate candidate loci further. In contrast, only a small number of autosomal windows were found to contain elevated signals for all three tests, so we focused on the two autosomal windows that had the most pronounced signals for the *BayPass* and IHS analysis: Chromosome 2 at ∼26.6 Mb and Chromosome 5 at ∼7.9 Mb (asterisks in Figure 5e).

**Figure 5:**
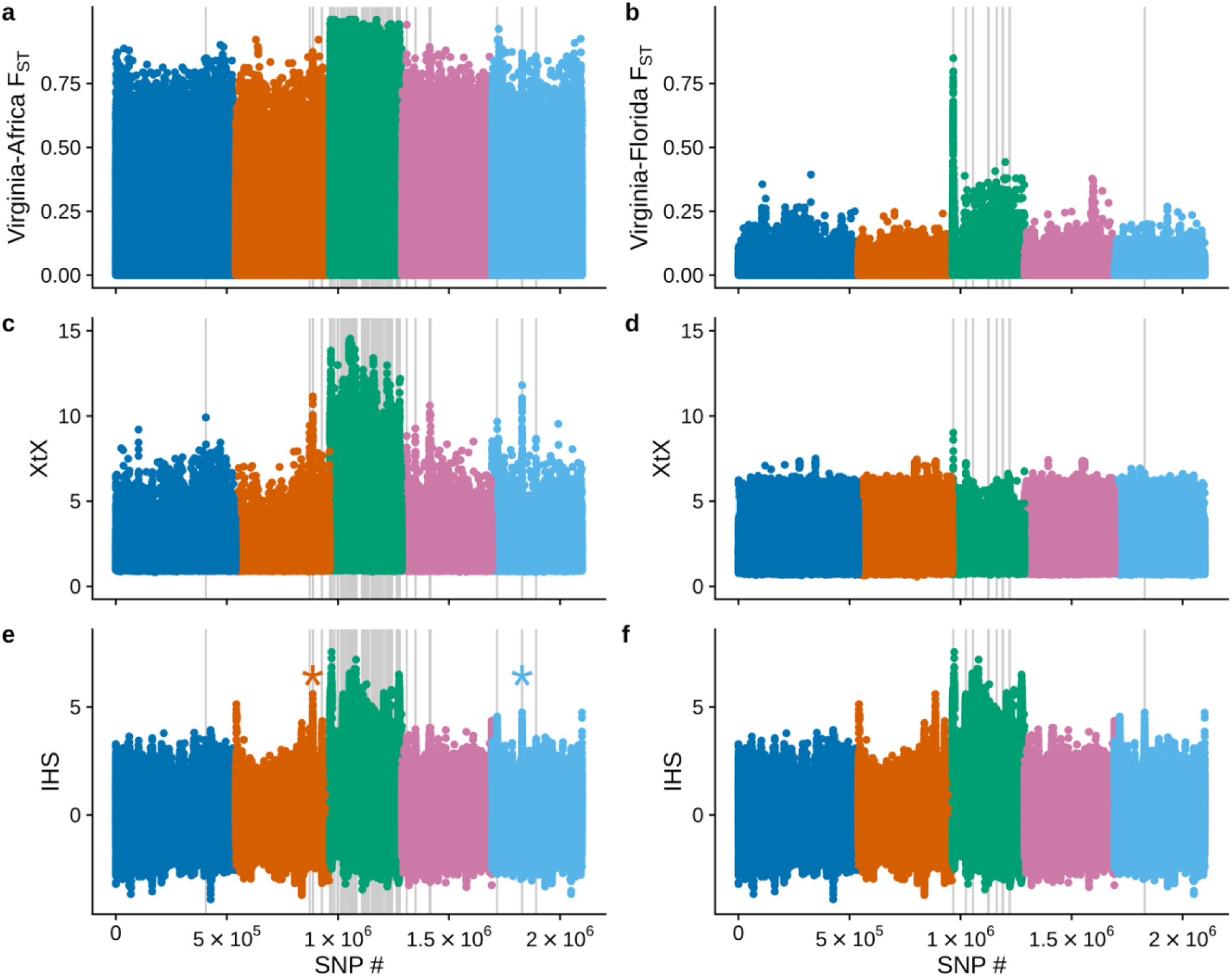
Signals of differentiation and selection in temperate Z. indianus populations. a-b) Genome-wide F_ST_ comparing individual flies sampled in Virginia (n= 175) to all flies sampled in Africa (a, n = 34) or Florida (b, n = 26); colors indicate chromosomes 1-5 in order. Only females were used for the X chromosome analysis (chromosome 3, green). c-d) BayPass XtX statistic for comparisons of females from Virginia and Africa (c) and Virginia and Florida (d). e-f) Integrated haplotype homozygosity score (IHS) testing for long, shared haplotypes in all flies collected in Virginia. The IHS datapoints in panels (e) and (f) are the same but presented twice to show alignments with each analysis above. Gray vertical lines indicate locations of 1 kb windows that contain F_ST_, XtX, and IHS scores in the top 1% genome-wide; windows were calculated separately for the Virginia-Africa and Virginia-Florida comparisons. Asterisks in panel (e) indicate the shared peaks on chromosomes 2 (orange) and 5 (light blue) discussed in the text and examined in Figure 6.

We examined the genes annotated in each of these outlier regions and found that both regions included genes in the cytochrome P450 pathway, which is involved in detoxifying endogenous and exogenous compounds including synthetic pesticides (Scott et al. 1998). The IHS peak on chromosome 2 is found near the *Z. indianus* homolog of *D. melanogaster cpr*, a cytochrome P450 reductase essential for electron transfer in the detoxification activities of cytochrome P450 genes (Hannemann et al. 2007). Decreased *cpr* expression has been linked to susceptibility to pesticides in a variety of insect pests and disease vectors (Lycett et al. 2006; Wang et al. 2016; He et al. 2020; Shi et al. 2021; Wang et al. 2025). A proline-to-leucine polymorphism in *cpr* has an XtX score of 7.63, is not found in Africa, and has a frequency of 0.57 in Virginia. Two other SNPs upstream of *cpr* also have large allele frequency differences between Virginia and Africa, offering interesting candidates for further study that might influence gene regulation.

The IHS peak on chromosome 5 is near a cluster of five cytochrome P450 genes. Although additional work will be required to resolve CYP gene orthologs and paralogs in *Z. indianus*, the cluster contains several *Cyp6* genes. One of the largest signals of selective sweeps in the *D. melanogaster* genome is found in *Cyp6g1* (Garud et al. 2015); alleles are *Cypg1* are associated with resistance to DDT and neonicotinoids (Daborn et al. 2001; Schmidt et al. 2010). Therefore, mutations in *Z. indianus CYP6* genes might also be involved in insecticide resistance. For example, in this region, missense variants in *Cyp6a9* and *Cyp317a1* have large allele frequency differences between Virginia and Africa; *Cyp317a1* was associated with permethrin resistance in the Drosophila Genetic Reference Panel (Battlay et al. 2018). *Z. indianus* is a substantial crop pest in Brazil, and several studies have indicated Brazilian *Z. indianus* populations may have evolved some degree of pesticide resistance in response to the use of organophosphorous compounds to protect fig crops (Galego and Carareto 2010; de Oliveira Rios et al. 2024). Although *Z. indianus* is not currently categorized as a pest in the United States (Animal and Plant Health Inspection Service, 2025) and is not an active target of management in eastern US orchards, many orchards use insecticides to control *D. suzukii* (Tait et al. 2021) and other arthropod pests. Controls for *D. suzukii* are often applied at the time of fruit ripening, which means that other species such as *Z. indianus* that are present during and after ripening may also be impacted by chemical control. Chemicals applied closer to the time of ripening may be more likely to remain on the overripe and decomposing fruit typically used as a breeding substrate for *Z. indianus.* Further investigation will be required to determine if mutations at these loci cause differences in pesticide resistance between African and North American flies. Insecticide resistance is a major challenge in the control of invasive insect pests (Siddiqui et al. 2023), and characterizing resistance may be important for future control of *Z. indianus*.

We confirmed the potential signal of selective sweeps by looking at haplotype structure in these two regions. Each region of high XtX score (Figure 6a-b) was characterized by a broad peak of extended haplotype homozygosity (EHH) specific to the allele found in invasive populations (Figure 6c-d), demonstrating that the candidate alleles have a slow decay of haplotype homozygosity. Visualization of the alleles found in each haplotype showed long haplotype blocks in North American populations that were not found in Africa (Figure 6e-f); SNPs within these blocks demonstrate high linkage disequilibrium in North American flies (Figure 6g-h). Additional analysis revealed these haplotypes were found in Colombia, suggesting they may have originated after *Z. indianus* arrived in the Western Hemisphere but may predate *Z. indianus’s* arrival in North America. In invasive copepods, haplotypes under selection in the invasive range are ancestral polymorphisms under balancing selection in the native range (Stern and Lee 2020). A similar situation was found for a balanced inversion polymorphism that fuels invasion in invasive crabs (Tepolt and Palumbi 2020; Tepolt et al. 2022). However, ancestral polymorphisms selected in the invaded range does not appear to be the case in *Z. indianus*, as the haplotypes from North America were not found in African flies. These novel haplotypes could be new mutations or derived due to hybridization/introgression from another species or divergent population; hybridization can be an important evolutionary force in invasive species (Ellstrand and Schierenbeck 2000; Fournier and Aron 2021). The *Zaprionus* genus shows signals of historic introgression among several species, though *Z. indianus* was not directly implicated in a previous analysis (Suvorov et al. 2022). Therefore, two major haplotypes not found in Africa have been selected in Florida and Virginia populations, though the source of these haplotypes remains to be determined.

**Figure 6:**
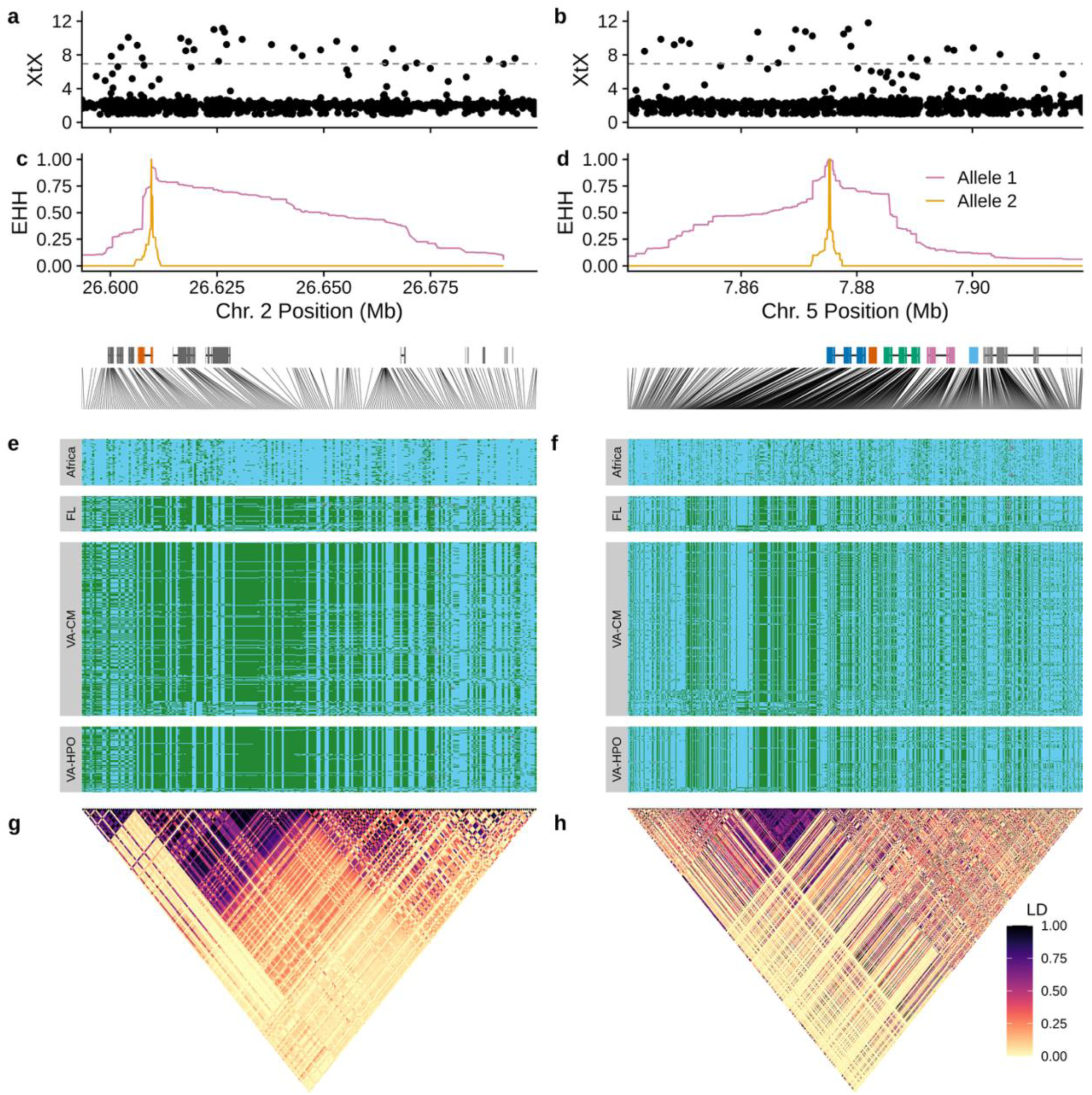
Haplotypes near potential pesticide resistance genes with signals of selection and differentiation. a-b) BayPass XtX statistics for individual SNPs comparing Africa and Virginia populations (see Figure 5a) for peaks on chromosomes 2 (a) and 5 (b). Horizontal lines indicate the 99.9% quantile of XtX scores from 10,000 simulated SNPs. c-d) Extended haplotype homozygosity for the SNP with highest IHS score at each locus (Chr. 2: 26,609,601 and Chr. 5: 7,875,359). The locations of annotated genes are shown below the x-axis. In panel (c) the gene highlighted in orange is Cytochrome P450 reductase (cpr). In (d) the genes highlighted in color are a cluster of five Cytochrome P450 genes (predicted as Cyp6a14, Cyp6a22, Cpy317a1, Cyp6a13, Cyp6a9 from left to right). Gray lines connect SNP order for panels below to physical locations on chromosomes. e-f) Haplotypes at each locus: each horizontal row shows genotypes for a single haploid chromosome phased with read-backed phasing. Blue indicates the allele more common in African populations and green is the other allele. Missing genotypes are shown in gray. Panel e shows 223 SNPs and panel f shows 599 SNPs. g-h) LD (R^2^) heatmaps for all SNPs for each haplotype demonstrate linkage in North American flies.

To explore population genetic signals around these potential regions of selection, we broadly grouped flies into three populations—Africa, Florida, and Virginia—and calculated population genetic statistics in 5 kb non-overlapping windows. F_ST_ between African and North American populations fell in the top 99% of all autosomal windows for both regions (Figure 7a-b), suggesting a high degree of genetic divergence. However, absolute genetic divergence (D_xy_) between African and North American populations was not especially elevated in these windows (Figure 7c-d). Nucleotide diversity (pi) was in the lowest 1% of all autosomal windows for North American samples (Figure 7e-f), as predicted for a sweep that results in shared haplotypes across many individuals. Tajima’s D was also negative and in the lowest 1% of all autosomal windows for North America, indicative of a recent sweep (Figure 7g-h). Tajima’s D was negative in Africa for these regions, but was it negative across the entire genome (Figure S15), likely due to combining genetically disparate subpopulations into a single population for this analysis, producing an excess of rare variants. Relative sequencing depth showed fluctuations in North American samples that were not seen in Africa (Figure 7h-i), suggesting the possibility of copy number variants may exist in both regions. Collectively, these tests provide additional evidence that both regions may have been subject to recent selective sweeps.

**Figure 7:**
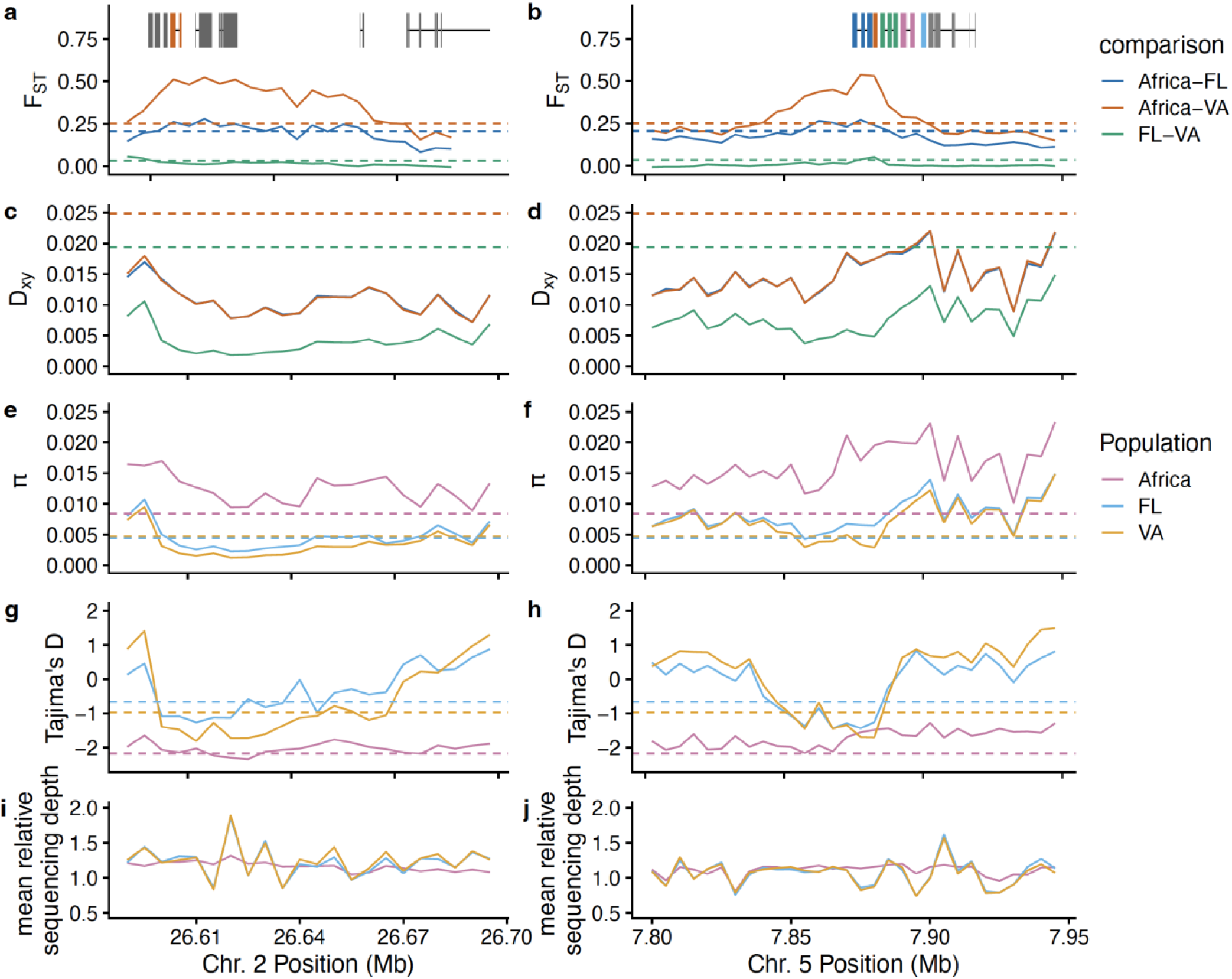
Population genetic statistics for the regions surrounding haplotypes on chromosomes 2 and 5. All statistics were calculated for 5 kb, non-overlapping windows. Panels on left focus on the chromosome 2 region containing cpr, and the panels on right focus on the chromosome 5 region containing several Cyp genes. Gene locations are shown at the top using same color scheme as in Figure 6. a-b) F_ST_ comparing combinations of flies from Africa, Virginia (both focal orchards combined) and Florida. c-d) Absolute nucleotide divergence (D_xy_) for the same comparisons. For a-d, horizontal lines indicate 99^th^ percentile of all autosomal windows. e-f) Nucleotide diversity (π) for each population. g-h) Tajima’s D for the three populations. For e-h, horizontal lines indicate bottom 1^st^ percentile of all autosomal windows. i-j) Average sequencing depth per window relative to the mean depth for the entire chromosome. Relative depths were averaged for all individuals in each population. See Figure S13 for whole-genome visualization of the same statistics.

For comparison, we also examined genetic diversity and population divergence across the genome (Figure S13) and observed that the X chromosome is an outlier in many regards. Divergence between Africa and North American samples is greater on the X chromosome (Figure S13A-B). As previously described (Comeault et al. 2020; Comeault et al. 2021), the X chromosome has reduced genetic diversity relative to autosomes, especially in invaded populations (Figure S13C). X chromosomes have smaller effective population sizes in species with XY sex determination systems and often experience more extreme loss of genetic diversity upon population contraction (Ellegren 2009). Tajima’s D is mostly positive in North American autosomes, indicative of a strong bottleneck in North American flies. However, Tajima’s D fluctuates between strongly positive and strongly negative in North American populations along the X chromosome (Figure S13D). These findings, combined with many regions with high haplotype homozygosity on the X chromosome (Figure 5e-f), suggest complex evolutionary dynamics on the X that warrant further investigation. Further global sampling and sequencing of X chromosomes with long reads to resolve inversion genotypes and CNVs may offer insight towards the potential role of X-linked variants in fueling the ongoing invasion of *Z. indianus*.

We lastly compared Virginia to Florida to test for possible recent adaptation within the invasive range of *Z. indianus*. Virginia has a temperate, seasonal climate with a relatively limited variety of cultivated produce, and southern Florida is subtropical with an abundance and diversity of fruits throughout the year. Other factors such as diseases, insecticide use, and competing species may also differ widely between locales. In the absence of genome-wide population structure, genomic regions differentiated between these locations are candidates for local adaptation. We observed elevated F_ST_ throughout much of the X chromosome, with a pronounced peak at 690 kb (Figure 5b); this peak was found near the most pronounced peaks in the XtX (Figure 5d) and IHS (Figure 5f) statistics. Examination of this region of the genome showed several long, shared haplotypes with high linkage in North American populations (Figure S14); in particular, one haplotype is nearly fixed in Virginia but is segregating in Florida. It is possible that this haplotype is associated with one of the structural variants on the X, and differences in inversion allele frequencies between Florida and Virginia could be driving these patterns. We examined population genetic signals in the region of highest F_ST_ (Figure S15) and determined that despite high F_ST_ from ∼550-650 kb (Figure S15A), D_xy_ between Florida and Virginia was not high in this region, nor were nucleotide diversity (Figure S15C) or Tajima’s D (Figure S15D) particularly low. In fact, immediately adjacent to the region of high FST was a region of nearly zero divergence between Florida and Virginia from ∼700-800 kb (Figure S15A-B). This region of low divergence had low nucleotide diversity (Figure S15C) and negative Tajima’s D (Figure S15D), indicating it could be a sweep in North America. However, it was immediately adjacent to a region of irregular sequencing depth (Figure S15E), suggesting the signals we observed could be confounded by read mapping artifacts.

Because the region of high F_ST_ is located near the end of the X chromosome and may be telomeric and/or centromeric, we more closely examined the normalized depth of sequencing coverage and presence of repetitive elements (Figure S16). This region contained a large repetitive element (Figure S16A), and sequencing depth was variable across populations (Figure S16B), suggesting structural variants such as duplications or deletions could be influencing mapping and population genetic statistics in this region. For example, at ∼610 kb, Florida samples had greater depth than Virginia, but from ∼625kb-650kb, Virginia samples had greater depth than Florida or Africa. All samples had low coverage near the repetitive element. These findings suggest copy number variation near repetitive elements at these loci might contribute to the Florida-Virginia divergence; whether the divergence is an artifact due to sequence misalignments or due to real copy number variation will require further analysis with long-read sequencing. The region of highest divergence between Florida and Viriginia contains ∼6 genes, including the gene *yin/opt1*, which is important for absorption of dietary peptides in *D. melanogaster* (Roman et al. 1998). If the Florida-Virginia divergence in this region is not a sequencing artifact, allelic differences within the invasive range in this gene could be involved in adaptation to new diets in new environments.

## Conclusions

In addition to posing economic, health, and environmental threats, invasive species also serve as outstanding models for studying rapid evolution in new environments. Here we report an improved genome assembly and annotation for *Z. indianus*, an introduced drosophilid that is thought to repeatedly recolonize temperate environments each year and is a potential crop pest. We use it for a preliminary assessment of potential rapid evolution and genetic variation in the early stages of invasion. We show that recolonization is likely a stochastic process resulting in different evolutionary dynamics in different years, even within a single orchard. This finding demonstrates broad sampling is important for invasive species that are repeatedly introduced or have multiple introduced populations that may undergo different evolutionary trajectories in different years or different locations. While some founding populations may be small, several population genetic patterns we observe could be explained by ongoing gene flow with the source population or between temperate populations following recolonization, suggesting gene flow that spreads and maintains favorable alleles could be an important component in *Z. indianus*’s widespread success, as it is for many invasive species (Díez-del-Molino et al. 2013; Medley et al. 2015; Arredondo et al. 2018). Demographic simulations and additional whole genome data will be required to better describe the recent histories of and potential gene flow between invasive populations and to infer colonization routes within North America.

Though we find limited population structure across space or time in introduced North American populations, we identified two selective sweeps in regions containing genes in the cytochrome P450 pathway, implicating pesticide resistance as a potential cause of recent sweeps in the invaded range. These haplotypes were not found in African samples, suggesting these alleles may have evolved in the past ∼25 years since *Z. indianus* invaded the Western Hemisphere, though we can’t rule out their origin in another undescribed population. Additional work will be required to characterize insecticide resistance in *Z. indianus* and compare resistance across alleles to test the hypothesis that these alleles are involved in a recent sweep involving pesticide resistance. We also find a region on the X chromosome that shows potential evidence of a selective sweep in temperate regions, potentially due to copy number variation, but the signal could be an artifact of genome assembly issues and sequencing near repetitive elements. Further investigation will be required to resolve the signals in this region. Studying how genetic variation in this region of the genome influences survival in temperate environments will be an important direction of future research. We additionally find that the X chromosome has an unusually complex evolutionary history in *Z. indianus*. It may have several segregating inversions and CNVs, has strong signatures of selection, and shows regions of high divergence both between African and North American populations and within North America. Specifically, long-read sequencing strategies will be important to understand likely inversions both on the X and throughout the *Z. indianus* genome that are common in the invaded range. Large inversions can link together adaptive alleles and are often important drivers of evolution in rapidly changing environments (Thompson and Jiggins 2014), so these regions will be important to track over larger spatial and temporal scales in future studies.

These results underscore the complexity of genetic dynamics during invasions and the need for further studies to explore the adaptive potential and ecological impacts of *Z. indianus* in its invasive range. *Z. indianus* provides a unique system in which we can study independent invasion events across multiple years and locations. One limitation of our study is sample size for each year and location: our ability to estimate allele frequencies or detect subtle changes in allele frequencies across time or space is limited. Sampling strategies that incorporate more individuals, such as pooled sequencing (Bergland et al. 2014; Kapun et al. 2021; Machado et al. 2021; Nunez et al. 2024), will be required to detect these more subtle changes, if they occur, and to understand how they may contribute to rapid adaptation to new environments. The recurrent nature of *Z. indianus* colonization may also offer insight towards the predictability of rapid evolution of invasive species.

## Data Availability

New individual sequencing data has been deposited in the SRA under project number # PRJNA991922. RNA sequencing from larval and pupal samples, and larval Hi-C data used for scaffolding are deposited under the same project number. The genome sequence has been deposited at DDBJ/ENA/GenBank under the accession JAUIZU000000000. The metadata for all sequencing samples (including date and location of collection); the annotation information for transcripts, proteins and repeats; and VCFs of SNPs and structural variants have been deposited to Dryad: https://doi.org/10.5061/dryad.q2bvq83v3. All code to reproduce analyses has been deposited to Zenodo via Dryad. All code for analysis is also available at: https://github.com/ericksonp/Z.indianus_individual_sequencing/tree/main

Temporary reviewer dryad link: http://datadryad.org/stash/share/6td1mtLMrbgLL6IgyaEtpmqK4cigV0Vl2Hhqw9Aspvo

## Acknowledgments

The authors acknowledge The University of Richmond’s High Performance Computer (https://data.richmond.edu/About-HPC-at-UR/index.html) for providing computational resources that contributed to the results reported herein. We particularly thank George Flanagin for technical support. Preliminary analyses were conducted using resources provided by Research Computing at The University of Virginia (https://rc.virginia.edu). We thank the owners and managers of Carter Mountain Orchard and Hanover Peach Orchard for graciously allowing us to collect flies on their properties. We thank Melinda Yang for helpful suggestions on demographic analyses.

## Author contributions

PAE, AB, AOB: conceptualization; PAE, AOB: funding; PAE, AG: investigation; PAE, AB, NP: resources; PAE, AS: formal analysis, PAE: methodology, visualization, writing-original draft; PAE, AOB: writing-reviewing and editing

## References

Alexander DH, Lange K. 2011. Enhancements to the ADMIXTURE algorithm for individual ancestry estimation. BMC Bioinformatics. 12(1):246. doi:10.1186/1471-2105-12-246. [accessed 2021 Jan 26]. https://doi.org/10.1186/1471-2105-12-246.

Allori Stazzonelli E, Funes CF, Corral Gonzalez MN, Gibilisco SM, Kirschbaum DS. 2023. Population fluctuation and infestation levels of Zaprionus indianus Gupta (Diptera: Drosophilidae) in berry crops of northwestern Argentina | International Society for Horticultural Science. Acta Horticultura.(1381). doi:10.17660/ActaHortic.2023.1381.19. [accessed 2024 Jun 19]. http://www.actahort.org/books/1381/1381_19.htm.

Altizer S, Ostfeld RS, Johnson PTJ, Kutz S, Harvell CD. 2013. Climate Change and Infectious Diseases: From Evidence to a Predictive Framework. Science. 341(6145):514–519. doi:10.1126/science.1239401.

Ananina G, Rohde C, David JR, Valente VLS, Klaczko LB. 2007. Inversion polymorphism and a new polytene chromosome map of Zaprionus indianus Gupta (1970) (Diptera: Drosophilidae). Genetica. 131(2):117–125. doi:10.1007/s10709-006-9121-6.

Animal and Plant Health Inspection Service. U.S. Regulated Plant Pest Table. [accessed 2025 Jun 9]. https://www.aphis.usda.gov/plant-imports/regulated-pest-list.

Araripe LO, Klaczko LB, Moreteau B, David JR. 2004. Male sterility thresholds in a tropical cosmopolitan drosophilid, Zaprionus indianus. Journal of Thermal Biology. 29(2):73–80. doi:10.1016/j.jtherbio.2003.11.006. [accessed 2019 Sep 5]. http://www.sciencedirect.com/science/article/pii/S0306456503000950.

Arredondo TM, Marchini GL, Cruzan MB. 2018. Evidence for human-mediated range expansion and gene flow in an invasive grass. Proceedings of the Royal Society B: Biological Sciences. 285(1882):20181125. doi:10.1098/rspb.2018.1125. [accessed 2024 Sep 5]. https://royalsocietypublishing.org/doi/full/10.1098/rspb.2018.1125.

Atsawawaranunt K, Ewart KM, Major RE, Johnson RN, Santure AW, Whibley A. 2023. Tracing the introduction of the invasive common myna using population genomics. Heredity. 131(1):56–67. doi:10.1038/s41437-023-00621-w. [accessed 2024 Jun 17]. https://www.nature.com/articles/s41437-023-00621-w.

Avalos A, Pan H, Li C, Acevedo-Gonzalez JP, Rendon G, Fields CJ, Brown PJ, Giray T, Robinson GE, Hudson ME, et al. 2017. A soft selective sweep during rapid evolution of gentle behaviour in an Africanized honeybee. Nat Commun. 8(1):1550. doi:10.1038/s41467-017-01800-0. [accessed 2024 Jun 25]. https://www.nature.com/articles/s41467-017-01800-0.

Bao Z, Eddy SR. 2002. Automated De Novo Identification of Repeat Sequence Families in Sequenced Genomes. Genome Res. 12(8):1269–1276. doi:10.1101/gr.88502. [accessed 2024 Feb 8]. https://genome.cshlp.org/content/12/8/1269.

Barrett CF, Corbett CW, Thixton-Nolan HL. 2023. A lack of population structure characterizes the invasive Lonicera japonica in West Virginia and across eastern North America1,2. tbot. 150(3):455–466. doi:10.3159/TORREY-D-23-00007.1. [accessed 2024 Jun 21]. https://bioone.org/journals/the-journal-of-the-torrey-botanical-society/volume-150/issue-3/TORREY-D-23-00007.1/A-lack-of-population-structure-characterizes-the-invasive-Lonicera-japonica/10.3159/TORREY-D-23-00007.1.full.

Barrett RDH, Laurent S, Mallarino R, Pfeifer SP, Xu CCY, Foll M, Wakamatsu K, Duke-Cohan JS, Jensen JD, Hoekstra HE. 2019. Linking a mutation to survival in wild mice. Science. 363(6426):499–504. doi:10.1126/science.aav3824.

Barrett SCH. 2015. Foundations of invasion genetics: the Baker and Stebbins legacy. Molecular Ecology. 24(9):1927–1941. doi:10.1111/mec.13014. [accessed 2019 Aug 29]. https://onlinelibrary.wiley.com/doi/abs/10.1111/mec.13014.

Battlay P, Leblanc PB, Green L, Garud NR, Schmidt JM, Fournier-Level A, Robin C. 2018. Structural Variants and Selective Sweep Foci Contribute to Insecticide Resistance in the Drosophila Genetic Reference Panel. G3 Genes|Genomes|Genetics. 8(11):3489–3497. doi:10.1534/g3.118.200619. [accessed 2021 Feb 9]. https://doi.org/10.1534/g3.118.200619.

Bellard C, Bertelsmeier C, Leadley P, Thuiller W, Courchamp F. 2012. Impacts of climate change on the future of biodiversity. Ecology Letters. 15(4):365–377. https://doi.org/10.1111/j.1461-0248.2011.01736.x.

Bemmels JB, Mikkelsen EK, Haddrath O, Colbourne RM, Robertson HA, Weir JT. 2021. Demographic decline and lineage-specific adaptations characterize New Zealand kiwi. Proc Biol Sci. 288(1965):20212362. doi:10.1098/rspb.2021.2362.

Bergland AO, Behrman EL, O’Brien KR, Schmidt PS, Petrov DA. 2014. Genomic Evidence of Rapid and Stable Adaptive Oscillations over Seasonal Time Scales in Drosophila. PLoS Genet. 10(11):e1004775. doi:10.1371/journal.pgen.1004775. [accessed 2015 Nov 2]. https://doi.org/10.1371/journal.pgen.1004775.

Bushnell B, Rood J, Singer E. 2017. BBMerge – Accurate paired shotgun read merging via overlap. PLOS ONE. 12(10):e0185056. doi:10.1371/journal.pone.0185056.

Cabanettes F, Klopp C. 2018. D-GENIES: dot plot large genomes in an interactive, efficient and simple way. PeerJ. 6:e4958. doi:10.7717/peerj.4958. [accessed 2024 Sep 18]. https://peerj.com/articles/4958.

Campos SRC, Rieger TT, Santos JF. 2007. Homology of polytene elements between Drosophila and Zaprionus determined by in situ hybridization in Zaprionus indianus. Genet Mol Res. 6(2):262–276.

Cantarel BL, Korf I, Robb SMC, Parra G, Ross E, Moore B, Holt C, Alvarado AS, Yandell M. 2008. MAKER: An easy-to-use annotation pipeline designed for emerging model organism genomes. Genome Res. 18(1):188–196. doi:10.1101/gr.6743907. [accessed 2024 Feb 8]. https://genome.cshlp.org/content/18/1/188.

Chan PP, Lowe TM. 2019. tRNAscan-SE: Searching for tRNA genes in genomic sequences. Methods Mol Biol. 1962:1–14. doi:10.1007/978-1-4939-9173-0_1. [accessed 2024 Feb 8]. https://www.ncbi.nlm.nih.gov/pmc/articles/PMC6768409/.

Chang CC, Chow CC, Tellier LC, Vattikuti S, Purcell SM, Lee JJ. 2015. Second-generation PLINK: rising to the challenge of larger and richer datasets. Gigascience. 4:7. doi:10.1186/s13742-015-0047-8.

Clements DR, Ditommaso A. 2011. Climate change and weed adaptation: can evolution of invasive plants lead to greater range expansion than forecasted? Weed Research. 51(3):227–240. https://doi.org/10.1111/j.1365-3180.2011.00850.x.

Comeault AA, Kautt AF, Matute DR. 2021. Genomic signatures of admixture and selection are shared among populations of Zaprionus indianus across the western hemisphere. Molecular Ecology. 30(23):6193–6210. doi:10.1111/mec.16066.

Comeault AA, Wang J, Tittes S, Isbell K, Ingley S, Hurlbert AH, Matute DR. 2020. Genetic Diversity and Thermal Performance in Invasive and Native Populations of African Fig Flies. Molecular Biology and Evolution. 37(7):1893–1906. doi:10.1093/molbev/msaa050.

Commar LS, Galego LG da C, Ceron CR, Carareto CMA. 2012. Taxonomic and evolutionary analysis of Zaprionus indianus and its colonization of Palearctic and Neotropical regions. Genet Mol Biol. 35(2):395–406. doi:10.1590/S1415-47572012000300003.

Daborn P, Boundy S, Yen J, Pittendrigh B, ffrench-Constant R. 2001. DDT resistance in Drosophila correlates with Cyp6g1 over-expression and confers cross-resistance to the neonicotinoid imidacloprid. Mol Gen Genomics. 266(4):556–563. doi:10.1007/s004380100531. [accessed 2025 May 29]. https://doi.org/10.1007/s004380100531.

Danecek P, Auton A, Abecasis G, Albers CA, Banks E, DePristo MA, Handsaker RE, Lunter G, Marth GT, Sherry ST, et al. 2011. The variant call format and VCFtools. Bioinformatics. 27(15):2156–2158. doi:10.1093/bioinformatics/btr330. [accessed 2024 Sep 17]. https://doi.org/10.1093/bioinformatics/btr330.

Danecek P, Bonfield JK, Liddle J, Marshall J, Ohan V, Pollard MO, Whitwham A, Keane T, McCarthy SA, Davies RM, et al. 2021. Twelve years of SAMtools and BCFtools. Gigascience. 10(2):giab008. doi:10.1093/gigascience/giab008.

Díez-del-Molino D, Carmona-Catot G, Araguas R-M, Vidal O, Sanz N, García-Berthou E, García-Marín J-L. 2013. Gene Flow and Maintenance of Genetic Diversity in Invasive Mosquitofish (Gambusia holbrooki). PLOS ONE. 8(12):e82501. doi:10.1371/journal.pone.0082501. [accessed 2024 Sep 5]. https://journals.plos.org/plosone/article?id=10.1371/journal.pone.0082501.

Dobin A, Davis CA, Schlesinger F, Drenkow J, Zaleski C, Jha S, Batut P, Chaisson M, Gingeras TR. 2013. STAR: ultrafast universal RNA-seq aligner. Bioinformatics. 29(1):15–21. doi:10.1093/bioinformatics/bts635. [accessed 2024 Feb 8]. https://www.ncbi.nlm.nih.gov/pmc/articles/PMC3530905/.

Dowle M, Srinivasan A. 2019. data.table: Extension of data.framè. https://CRAN.R-project.org/package=data.table.

Ellegren H. 2009. The different levels of genetic diversity in sex chromosomes and autosomes. Trends in Genetics. 25(6):278–284. doi:10.1016/j.tig.2009.04.005. [accessed 2024 Jun 24]. https://www.sciencedirect.com/science/article/pii/S0168952509000900.

Ellstrand NC, Schierenbeck KA. 2000. Hybridization as a stimulus for the evolution of invasiveness in plants? Proceedings of the National Academy of Sciences. 97(13):7043–7050. doi:10.1073/pnas.97.13.7043. [accessed 2024 Jun 21]. https://www.pnas.org/doi/abs/10.1073/pnas.97.13.7043.

Erickson PA, Weller CA, Song DY, Bangerter AS, Schmidt P, Bergland AO. 2020. Unique genetic signatures of local adaptation over space and time for diapause, an ecologically relevant complex trait, in Drosophila melanogaster. PLOS Genetics. 16(11):e1009110. doi:10.1371/journal.pgen.1009110.

Estoup A, Ravigné V, Hufbauer R, Vitalis R, Gautier M, Facon B. 2016. Is There a Genetic Paradox of Biological Invasion? Annual Review of Ecology, Evolution, and Systematics. 47(Volume 47, 2016):51–72. doi:10.1146/annurev-ecolsys-121415-032116. [accessed 2025 Jun 11]. https://www.annualreviews.org/content/journals/10.1146/annurev-ecolsys-121415-032116.

Fang Z, Pyhäjärvi T, Weber AL, Dawe RK, Glaubitz JC, González J de JS, Ross-Ibarra C, Doebley J, Morrell PL, Ross-Ibarra J. 2012. Megabase-Scale Inversion Polymorphism in the Wild Ancestor of Maize. Genetics. 191(3):883–894. doi:10.1534/genetics.112.138578. [accessed 2024 Jun 24]. https://doi.org/10.1534/genetics.112.138578.

Felsenstein J. 2005. PHYLIP (Phylogeny Inference Package) version 3.6. https://phylipweb.github.io/phylip/.

Feron R, Waterhouse RM. 2022. Assessing species coverage and assembly quality of rapidly accumulating sequenced genomes. GigaScience. 11:giac006. doi:10.1093/gigascience/giac006. [accessed 2024 Jun 20]. https://doi.org/10.1093/gigascience/giac006.

Flynn JM, Hubley R, Goubert C, Rosen J, Clark AG, Feschotte C, Smit AF. 2020. RepeatModeler2 for automated genomic discovery of transposable element families. Proceedings of the National Academy of Sciences. 117(17):9451–9457. doi:10.1073/pnas.1921046117. [accessed 2024 Feb 8]. https://www.pnas.org/doi/full/10.1073/pnas.1921046117.

Fournier D, Aron S. 2021. Hybridization and invasiveness in social insects — The good, the bad and the hybrid. Current Opinion in Insect Science. 46:1–9. doi:10.1016/j.cois.2020.12.004. [accessed 2024 Jun 21]. https://www.sciencedirect.com/science/article/pii/S2214574521000018.

Friedline CJ, Faske TM, Lind BM, Hobson EM, Parry D, Dyer RJ, Johnson DM, Thompson LM, Grayson KL, Eckert AJ. 2019. Evolutionary genomics of gypsy moth populations sampled along a latitudinal gradient. Molecular Ecology. 0(0). doi:10.1111/mec.15069. [accessed 2019 Jun 6]. https://onlinelibrary.wiley.com/doi/abs/10.1111/mec.15069.

Galego LG da C, Carareto CMA. 2010. Variation at the Est3 locus and adaptability to organophosphorous compounds in Zaprionus indianus populations. Entomologia Experimentalis et Applicata. 134(1):97–105. doi:10.1111/j.1570-7458.2009.00941.x. [accessed 2025 May 29]. https://onlinelibrary.wiley.com/doi/abs/10.1111/j.1570-7458.2009.00941.x.

Galludo M, Canals J, Pineda-Cirera L, Esteve C, Rosselló M, Balanyà J, Arenas C, Mestres F. 2018. Climatic adaptation of chromosomal inversions in Drosophila subobscura. Genetica. 146(4):433–441. doi:10.1007/s10709-018-0035-x. [accessed 2024 Jun 20]. https://doi.org/10.1007/s10709-018-0035-x.

García-Escudero CA, Tsigenopoulos CS, Manousaki T, Tsakogiannis A, Marbà N, Vizzini S, Duarte CM, Apostolaki ET. 2023. Population genomics unveils the century-old invasion of the Seagrass Halophila stipulacea in the Mediterranean Sea. Mar Biol. 171(2):40. doi:10.1007/s00227-023-04361-7. [accessed 2024 Jun 17]. https://doi.org/10.1007/s00227-023-04361-7.

Garnier S. 2018. viridis: Default Color Maps from “matplotlib.” https://CRAN.R-project.org/package=viridis.

Garud NR, Messer PW, Buzbas EO, Petrov DA. 2015. Recent Selective Sweeps in North American Drosophila melanogaster Show Signatures of Soft Sweeps. PLOS Genetics. 11(2):e1005004. doi:10.1371/journal.pgen.1005004. [accessed 2021 Jan 21]. https://journals.plos.org/plosgenetics/article?id=10.1371/journal.pgen.1005004.

Gautier M. 2015. Genome-Wide Scan for Adaptive Divergence and Association with Population-Specific Covariates. Genetics. 201(4):1555–1579. doi:10.1534/genetics.115.181453. [accessed 2025 May 27]. https://doi.org/10.1534/genetics.115.181453.

Gautier M, Vitalis R. 2012. rehh: an R package to detect footprints of selection in genome-wide SNP data from haplotype structure. Bioinformatics. 28(8):1176–1177. doi:10.1093/bioinformatics/bts115.

Gleason JM, Roy PR, Everman ER, Gleason TC, Morgan TJ. 2019. Phenology of Drosophila species across a temperate growing season and implications for behavior. PLOS ONE. 14(5):e0216601. doi:10.1371/journal.pone.0216601.

Good BH, McDonald MJ, Barrick JE, Lenski RE, Desai MM. 2017. The dynamics of molecular evolution over 60,000 generations. Nature. 551(7678):45–50. doi:10.1038/nature24287.

Gupta JP. 1970. Description of a new species of Phorticella zaprionus (Drosophilidae) from India. Proceedings of the Indian National Science Academy. 36B(1):62–70.

Gupta JP, Kumar A. 1987. Cytogenetics of Zaprionus indianus Gupta (Diptera: Drosophilidae): Nucleolar organizer regions, mitotic and polytene chromosomes and inversion polymorphism. Genetica. 74(1):19–25. doi:10.1007/BF00055090. [accessed 2023 Jul 13]. https://doi.org/10.1007/BF00055090.

Hancock AM, Brachi B, Faure N, Horton MW, Jarymowycz LB, Sperone FG, Toomajian C, Roux F, Bergelson J. 2011. Adaptation to Climate Across the Arabidopsis thaliana Genome. Science. 334(6052):83–86. doi:10.1126/science.1209244.

Hannemann F, Bichet A, Ewen KM, Bernhardt R. 2007. Cytochrome P450 systems—biological variations of electron transport chains. Biochimica et Biophysica Acta (BBA) - General Subjects. 1770(3):330–344. doi:10.1016/j.bbagen.2006.07.017. [accessed 2025 May 29]. https://www.sciencedirect.com/science/article/pii/S0304416506002133.

He C, Liang J, Liu S, Zeng Y, Wang S, Wu Q, Xie W, Zhang Y. 2020. Molecular characterization of an NADPH cytochrome P450 reductase from *Bemisia tabaci* Q: Potential involvement in susceptibility to imidacloprid. Pesticide Biochemistry and Physiology. 162:29–35. doi:10.1016/j.pestbp.2019.07.018. [accessed 2025 May 29]. https://www.sciencedirect.com/science/article/pii/S0048357519304420.

Hoberg EP, Brooks DR. 2015. Evolution in action: climate change, biodiversity dynamics and emerging infectious disease. Philosophical Transactions of the Royal Society B: Biological Sciences. 370(1665):20130553. doi:10.1098/rstb.2013.0553.

Holle SG, Tran AK, Burkness EC, Ebbenga DN, Hutchison WD. 2018. First Detections of Zaprionus indianus (Diptera: Drosophilidae) in Minnesota. ents. 54(1):99–102. doi:10.18474/JES18-22.

Huang K, Andrew RL, Owens GL, Ostevik KL, Rieseberg LH. 2020. Multiple chromosomal inversions contribute to adaptive divergence of a dune sunflower ecotype. Molecular Ecology. 29(14):2535–2549. doi:10.1111/mec.15428. [accessed 2024 Jun 20]. https://onlinelibrary.wiley.com/doi/abs/10.1111/mec.15428.

Johnson MS, Gopalakrishnan S, Goyal J, Dillingham ME, Bakerlee CW, Humphrey PT, Jagdish T, Jerison ER, Kosheleva K, Lawrence KR, et al. 2021. Phenotypic and molecular evolution across 10,000 generations in laboratory budding yeast populations. Verstrepen KJ, Wittkopp PJ, Verstrepen KJ, Hodgins-Davis A, editors. eLife. 10:e63910. doi:10.7554/eLife.63910.

Jombart T. 2008. adegenet: a R package for the multivariate analysis of genetic markers. Bioinformatics. 24(11):1403–1405. doi:10.1093/bioinformatics/btn129. [accessed 2025 May 27]. https://doi.org/10.1093/bioinformatics/btn129.

Jombart T, Ahmed I. 2011. adegenet 1.3-1: new tools for the analysis of genome-wide SNP data. Bioinformatics. 27(21):3070–3071. doi:10.1093/bioinformatics/btr521. [accessed 2025 May 27]. https://doi.org/10.1093/bioinformatics/btr521.

Jones MR, Mills LS, Alves PC, Callahan CM, Alves JM, Lafferty DJR, Jiggins FM, Jensen JD, Melo-Ferreira J, Good JM. 2018. Adaptive introgression underlies polymorphic seasonal camouflage in snowshoe hares. Science. 360(6395):1355–1358. doi:10.1126/science.aar5273.

Joron M, Frezal L, Jones RT, Chamberlain NL, Lee SF, Haag CR, Whibley A, Becuwe M, Baxter SW, Ferguson L, et al. 2011. Chromosomal rearrangements maintain a polymorphic supergene controlling butterfly mimicry. Nature. 477(7363):203–206. doi:10.1038/nature10341. [accessed 2015 Sep 14]. http://www.nature.com/nature/journal/v477/n7363/full/nature10341.html.

Joshi NK, Biddinger DJ, Demchak K, Deppen A. 2014. First report of Zaprionus indianus (Diptera: Drosophilidae) in commercial fruits and vegetables in Pennsylvania. J Insect Sci. 14:259. doi:10.1093/jisesa/ieu121.

Kapun M, Nunez JCB, Bogaerts-Márquez M, Murga-Moreno J, Paris M, Outten J, Coronado-Zamora M, Tern C, Rota-Stabelli O, Guerreiro MPG, et al. 2021. Drosophila Evolution over Space and Time (DEST): A New Population Genomics Resource. Molecular Biology and Evolution. 38(12):5782–5805. doi:10.1093/molbev/msab259. [accessed 2025 Jun 13]. https://doi.org/10.1093/molbev/msab259.

Keller O, Kollmar M, Stanke M, Waack S. 2011. A novel hybrid gene prediction method employing protein multiple sequence alignments. Bioinformatics. 27(6):757–763. doi:10.1093/bioinformatics/btr010. [accessed 2024 Feb 8]. https://doi.org/10.1093/bioinformatics/btr010.

Kim BY, Wang JR, Miller DE, Barmina O, Delaney E, Thompson A, Comeault AA, Peede D, D’Agostino ER, Pelaez J, et al. 2021. Highly contiguous assemblies of 101 drosophilid genomes. Coop G, Wittkopp PJ, Sackton TB, editors. eLife. 10:e66405. doi:10.7554/eLife.66405. [accessed 2023 Jul 13]. https://doi.org/10.7554/eLife.66405.

Koch JB, Dupuis JR, Jardeleza M-K, Ouedraogo N, Geib SM, Follett PA, Price DK. 2020. Population genomic and phenotype diversity of invasive Drosophila suzukii in Hawai‘i. Biol Invasions. 22(5):1753–1770. doi:10.1007/s10530-020-02217-5. [accessed 2024 Jun 17]. https://doi.org/10.1007/s10530-020-02217-5.

Kohlmeier Pinar, Kohlmeier Philip. 2025. A non-overwintering urban population of the African fig fly (Diptera: Drosophilidae) impacts the reproductive output of locally adapted fruit flies. Florida Entomologist. 108(1). doi:10.1515/flaent-2024-0064. [accessed 2025 Mar 29]. https://www.degruyter.com/document/doi/10.1515/flaent-2024-0064/html?srsltid=AfmBOoqia32bki8F2DqWqVQeWln4SQ5boCJG7oVXUNkgD7qLXESk_noP.

Korf I. 2004. Gene finding in novel genomes. BMC Bioinformatics. 5(1):59. doi:10.1186/1471-2105-5-59. [accessed 2024 Feb 8]. https://doi.org/10.1186/1471-2105-5-59.

Korneliussen TS, Albrechtsen A, Nielsen R. 2014. ANGSD: Analysis of Next Generation Sequencing Data. BMC Bioinformatics. 15(1):356. doi:10.1186/s12859-014-0356-4. [accessed 2023 Jul 13]. https://doi.org/10.1186/s12859-014-0356-4.

Korunes KL, Samuk K. pixy: Unbiased estimation of nucleotide diversity and divergence in the presence of missing data. Molecular Ecology Resources. n/a(n/a). 10.1111/1755-0998.13326. [accessed 2021 Feb 12]. https://onlinelibrary.wiley.com/doi/abs/10.1111/1755-0998.13326.

Kremmer L, David J, Borowiec N, Thaon M, Ris N, Poirie M, Gatti J-L. 2017. The African fig fly Zaprionus indianus: a new invasive pest in France? Bulletin of Insectology. 70(1):57–62.

Küpper C, Stocks M, Risse JE, dos Remedios N, Farrell LL, McRae SB, Morgan TC, Karlionova N, Pinchuk P, Verkuil YI, et al. 2016. A supergene determines highly divergent male reproductive morphs in the ruff. Nat Genet. 48(1):79–83. doi:10.1038/ng.3443. [accessed 2016 Mar 14]. http://www.nature.com/ng/journal/v48/n1/full/ng.3443.html.

Leão BFD, Tldon R. 2004. Newly invading species exploiting native host-plants: the case of the African Zaprionus indianus (Gupta) in the Brazilian Cerrado (Diptera, Drosophilidae). Annales de la Société entomologique de France (NS). 40(3–4):285–290. doi:10.1080/00379271.2004.10697427.

Lee YW, Fishman L, Kelly JK, Willis JH. 2016. A Segregating Inversion Generates Fitness Variation in Yellow Monkeyflower (Mimulus guttatus). Genetics. 202(4):1473–1484. doi:10.1534/genetics.115.183566. [accessed 2016 Apr 22]. http://www.genetics.org/content/202/4/1473.

Li H, Durbin R. 2009. Fast and accurate short read alignment with Burrows-Wheeler transform. Bioinformatics. 25(14):1754–1760. doi:10.1093/bioinformatics/btp324.

Li H, Handsaker B, Wysoker A, Fennell T, Ruan J, Homer N, Marth G, Abecasis G, Durbin R, 1000 Genome Project Data Processing Subgroup. 2009. The Sequence Alignment/Map format and SAMtools. Bioinformatics. 25(16):2078–2079. doi:10.1093/bioinformatics/btp352. [accessed 2022 Dec 9]. https://doi.org/10.1093/bioinformatics/btp352.

Li H, Peng Y, Wang Y, Summerhays B, Shu X, Vasquez Y, Vansant H, Grenier C, Gonzalez N, Kansagra K, et al. 2023. Global patterns of genomic and phenotypic variation in the invasive harlequin ladybird. BMC Biol. 21(1):141. doi:10.1186/s12915-023-01638-7. [accessed 2024 Jun 17]. https://doi.org/10.1186/s12915-023-01638-7.

Li H, Ralph P. 2019. Local PCA Shows How the Effect of Population Structure Differs Along the Genome. Genetics. 211(1):289–304. doi:10.1534/genetics.118.301747. [accessed 2024 May 14]. https://www.ncbi.nlm.nih.gov/pmc/articles/PMC6325702/.

Linck E, Battey CJ. 2019. Minor allele frequency thresholds strongly affect population structure inference with genomic data sets. Molecular Ecology Resources. 19(3):639–647. doi:10.1111/1755-0998.12995. [accessed 2025 Jun 3]. https://onlinelibrary.wiley.com/doi/abs/10.1111/1755-0998.12995.

Linde K van der, Steck GJ, Hibbard K, Birdsley JS, Alonso LM, Houle D. 2006. FIRST RECORDS OF ZAPRIONUS INDIANUS (DIPTERA: DROSOPHILIDAE), A PEST SPECIES ON COMMERCIAL FRUITS FROM PANAMA AND THE UNITED STATES OF AMERICA. flen. 89(3):402–404. doi:10.1653/0015-4040(2006)89[402:FROZID]2.0.CO;2.

Lovell JT, MacQueen AH, Mamidi S, Bonnette J, Jenkins J, Napier JD, Sreedasyam A, Healey A, Session A, Shu S, et al. 2021 Jan 27. Genomic mechanisms of climate adaptation in polyploid bioenergy switchgrass. Nature.:1–7. doi:10.1038/s41586-020-03127-1.

Lycett GJ, McLaughlin LA, Ranson H, Hemingway J, Kafatos FC, Loukeris TG, Paine MJI. 2006. Anopheles gambiae P450 reductase is highly expressed in oenocytes and in vivo knockdown increases permethrin susceptibility. Insect Molecular Biology. 15(3):321–327. doi:10.1111/j.1365-2583.2006.00647.x. [accessed 2025 May 26]. https://onlinelibrary.wiley.com/doi/abs/10.1111/j.1365-2583.2006.00647.x.

Ma L, Cao L-J, Hoffmann AA, Gong Y-J, Chen J-C, Chen H-S, Wang X-B, Zeng A-P, Wei S-J, Zhou Z-S. 2020. Rapid and strong population genetic differentiation and genomic signatures of climatic adaptation in an invasive mealybug. Diversity and Distributions. 26(5):610–622. 10.1111/ddi.13053. [accessed 2021 Jan 5]. https://onlinelibrary.wiley.com/doi/abs/10.1111/ddi.13053.

Ma L-J, Cao L-J, Chen J-C, Tang M-Q, Song W, Yang F-Y, Shen X-J, Ren Y-J, Yang Q, Li H, et al. 2024. Rapid and Repeated Climate Adaptation Involving Chromosome Inversions following Invasion of an Insect. Molecular Biology and Evolution. 41(3):msae044. doi:10.1093/molbev/msae044. [accessed 2024 Jun 17]. https://doi.org/10.1093/molbev/msae044.

Machado HE, Bergland AO, Taylor R, Tilk S, Behrman E, Dyer K, Fabian DK, Flatt T, González J, Karasov TL, et al. 2021. Broad geographic sampling reveals the shared basis and environmental correlates of seasonal adaptation in Drosophila. Nordborg M, Wittkopp PJ, Nordborg M, editors. eLife. 10:e67577. doi:10.7554/eLife.67577. [accessed 2024 Jun 20]. https://doi.org/10.7554/eLife.67577.

Manni M, Berkeley MR, Seppey M, Simão FA, Zdobnov EM. 2021. BUSCO Update: Novel and Streamlined Workflows along with Broader and Deeper Phylogenetic Coverage for Scoring of Eukaryotic, Prokaryotic, and Viral Genomes. Molecular Biology and Evolution. 38(10):4647– 4654. doi:10.1093/molbev/msab199. [accessed 2024 Sep 19]. https://doi.org/10.1093/molbev/msab199.

Markow TA, Hanna G, Riesgo-Escovar JR, Tellez-Garcia AA, Richmond MP, Nazario-Yepiz NO, Laclette MRL, Carpinteyro-Ponce J, Pfeiler E. 2014. Population genetics and recent colonization history of the invasive drosophilid Zaprionus indianus in Mexico and Central America. Biol Invasions. 16(11):2427–2434. doi:10.1007/s10530-014-0674-5.

da Mata RA, Tidon R, Côrtes LG, De Marco P, Diniz-Filho JAF. 2010. Invasive and flexible: niche shift in the drosophilid Zaprionus indianus (Insecta, Diptera). Biol Invasions. 12(5):1231– 1241. doi:10.1007/s10530-009-9542-0.

Matheson P, McGaughran A. 2022. Genomic data is missing for many highly invasive species, restricting our preparedness for escalating incursion rates. Sci Rep. 12(1):13987. doi:10.1038/s41598-022-17937-y. [accessed 2024 Jun 17]. https://www.nature.com/articles/s41598-022-17937-y.

McGaughran A, Dhami MK, Parvizi E, Vaughan AL, Gleeson DM, Hodgins KA, Rollins LA, Tepolt CK, Turner KG, Atsawawaranunt K, et al. 2024. Genomic Tools in Biological Invasions: Current State and Future Frontiers. Genome Biology and Evolution. 16(1):evad230. doi:10.1093/gbe/evad230. [accessed 2024 Jun 17]. https://doi.org/10.1093/gbe/evad230.

McKenna A, Hanna M, Banks E, Sivachenko A, Cibulskis K, Kernytsky A, Garimella K, Altshuler D, Gabriel S, Daly M, et al. 2010. The Genome Analysis Toolkit: a MapReduce framework for analyzing next-generation DNA sequencing data. Genome Res. 20(9):1297– 1303. doi:10.1101/gr.107524.110.

Medley KA, Jenkins DG, Hoffman EA. 2015. Human-aided and natural dispersal drive gene flow across the range of an invasive mosquito. Molecular Ecology. 24(2):284–295. doi:10.1111/mec.12925. [accessed 2024 Sep 5]. https://onlinelibrary.wiley.com/doi/abs/10.1111/mec.12925.

Messer PW, Petrov DA. 2013. Population genomics of rapid adaptation by soft selective sweeps. Trends in Ecology & Evolution. 28(11):659–669. doi:10.1016/j.tree.2013.08.003. [accessed 2024 Jun 20]. https://www.cell.com/trends/ecology-evolution/abstract/S0169-5347(13)00207-3.

Microsoft, Weston S. 2017. foreach: Provides Foreach Looping Construct for R. https://CRAN.R-project.org/package=foreach.

Nava DE, Nascimento AM, Stein CP, Haddad ML, Bento JMS, Parra JRP. 2007. Biology, thermal requirements, and estimation of the number of generations of Zaprionus indianus (Diptera: Drosopholidae) for the main fig producing regions of Brazil. flen. 90(3):495–501. doi:10.1653/0015-4040(2007)90[495:BTRAEO]2.0.CO;2.

Nguyen Ba AN, Cvijović I, Rojas Echenique JI, Lawrence KR, Rego-Costa A, Liu X, Levy SF, Desai MM. 2019. High-resolution lineage tracking reveals travelling wave of adaptation in laboratory yeast. Nature. 575(7783):494–499. doi:10.1038/s41586-019-1749-3.

Nowling RJ, Manke KR, Emrich SJ. 2020. Detecting inversions with PCA in the presence of population structure. PLOS ONE. 15(10):e0240429. doi:10.1371/journal.pone.0240429. [accessed 2024 Jun 20]. https://journals.plos.org/plosone/article?id=10.1371/journal.pone.0240429.

Nunez JCB, Lenhart BA, Bangerter A, Murray CS, Mazzeo GR, Yu Y, Nystrom TL, Tern C, Erickson PA, Bergland AO. 2024. A cosmopolitan inversion facilitates seasonal adaptation in overwintering Drosophila. Genetics. 226(2):iyad207. doi:10.1093/genetics/iyad207. [accessed 2024 Jun 20]. https://doi.org/10.1093/genetics/iyad207.

Oerke E-C. 2006. Crop losses to pests. The Journal of Agricultural Science. 144(1):31–43. doi:10.1017/S0021859605005708.

Oliveira CM, Auad AM, Mendes SM, Frizzas MR. 2013. Economic impact of exotic insect pests in Brazilian agriculture. Journal of Applied Entomology. 137(1–2):1–15. 10.1111/jen.12018.

de Oliveira Rios J, Costa SC, da Conceição Galego LG. 2024. Geographical and sex-specific effects of malathion insecticide selection in African fig fly Zaprionus indianus Gupta, 1970 (Diptera: Drosophilidae). Journal of Applied Entomology. 148(3):312–320. doi:10.1111/jen.13212. [accessed 2025 May 29]. https://onlinelibrary.wiley.com/doi/abs/10.1111/jen.13212.

Parchami-Araghi M, Gilasian E, Keyhanian A. 2015. Olive infestation with Zaprionus indianus Gupta (Dip.: Drosophilidae) in northern Iran: a new host record and threat to world olive production. Drosophila Information Service. 98:60–61.

Parvizi E, Dhami MK, Yan J, McGaughran A. 2023. Population genomic insights into invasion success in a polyphagous agricultural pest, Halyomorpha halys. Molecular Ecology. 32(1):138–151. doi:10.1111/mec.16740. [accessed 2024 Jun 17]. https://onlinelibrary.wiley.com/doi/abs/10.1111/mec.16740.

Patterson M, Marschall T, Pisanti N, van Iersel L, Stougie L, Klau GW, Schönhuth A. 2015. WhatsHap: Weighted Haplotype Assembly for Future-Generation Sequencing Reads. J Comput Biol. 22(6):498–509. doi:10.1089/cmb.2014.0157.

Patton AH, Margres MJ, Stahlke AR, Hendricks S, Lewallen K, Hamede RK, Ruiz-Aravena M, Ryder O, McCallum HI, Jones ME, et al. 2019. Contemporary Demographic Reconstruction Methods Are Robust to Genome Assembly Quality: A Case Study in Tasmanian Devils. Molecular Biology and Evolution. 36(12):2906–2921. doi:10.1093/molbev/msz191. [accessed 2021 Jan 13]. https://doi.org/10.1093/molbev/msz191.

Pedersen BS, Layer R, Quinlan AR. 2020. smoove: structural-variant calling and genotyping with existing tools.

Pélissié B, Chen YH, Cohen ZP, Crossley MS, Hawthorne DJ, Izzo V, Schoville SD. 2022. Genome Resequencing Reveals Rapid, Repeated Evolution in the Colorado Potato Beetle. Molecular Biology and Evolution. 39(2):msac016. doi:10.1093/molbev/msac016. [accessed 2024 Jun 17]. https://doi.org/10.1093/molbev/msac016.

Pfeiffer DG, Shrader ME, Wahls JCE, Willbrand BN, Sandum I, van der Linde K, Laub CA, Mays RS, Day ER. 2019. African Fig Fly (Diptera: Drosophilidae): Biology, Expansion of Geographic Range, and Its Potential Status as a Soft Fruit Pest. J Integr Pest Manag. 10(1). doi:10.1093/jipm/pmz018. [accessed 2019 Aug 15]. https://academic.oup.com/jipm/article/10/1/20/5514212.

Pickrell JK, Pritchard JK. 2012. Inference of Population Splits and Mixtures from Genome-Wide Allele Frequency Data. PLOS Genetics. 8(11):e1002967. doi:10.1371/journal.pgen.1002967. [accessed 2025 May 29]. https://journals.plos.org/plosgenetics/article?id=10.1371/journal.pgen.1002967.

Picq S, Wu Y, Martemyanov VV, Pouliot E, Pfister SE, Hamelin R, Cusson M. 2023. Range-wide population genomics of the spongy moth, Lymantria dispar (Erebidae): Implications for biosurveillance, subspecies classification and phylogeography of a destructive moth. Evolutionary Applications. 16(3):638–656. doi:10.1111/eva.13522. [accessed 2024 Jun 20]. https://onlinelibrary.wiley.com/doi/abs/10.1111/eva.13522.

Platts PJ, Mason SC, Palmer G, Hill JK, Oliver TH, Powney GD, Fox R, Thomas CD. 2019. Habitat availability explains variation in climate-driven range shifts across multiple taxonomic groups. Scientific Reports. 9(1):15039. doi:10.1038/s41598-019-51582-2.

Price AL, Jones NC, Pevzner PA. 2005. De novo identification of repeat families in large genomes. Bioinformatics. 21(suppl_1):i351–i358. doi:10.1093/bioinformatics/bti1018. [accessed 2024 Feb 8]. https://doi.org/10.1093/bioinformatics/bti1018.

Purcell S, Neale B, Todd-Brown K, Thomas L, Ferreira MAR, Bender D, Maller J, Sklar P, de Bakker PIW, Daly MJ, et al. 2007. PLINK: a tool set for whole-genome association and population-based linkage analyses. Am J Hum Genet. 81(3):559–575. doi:10.1086/519795.

Putnam NH, O’Connell BL, Stites JC, Rice BJ, Blanchette M, Calef R, Troll CJ, Fields A, Hartley PD, Sugnet CW, et al. 2016. Chromosome-scale shotgun assembly using an in vitro method for long-range linkage. Genome Res. 26(3):342–350. doi:10.1101/gr.193474.115. [accessed 2023 Jul 13]. https://www.ncbi.nlm.nih.gov/pmc/articles/PMC4772016/.

R Core Team. R: A language and environment for statistical computing. http://www.R-project.org/.

Rakes LM, Delamont M, Cole C, Yates JA, Blevins LJ, Hassan FN, Bergland AO, Erickson PA. 2023. A small survey of introduced Zaprionus indianus (Diptera: Drosophilidae) in orchards of the eastern United States. Journal of Insect Science. 23(5):21. doi:10.1093/jisesa/iead092. [accessed 2024 Feb 8]. https://doi.org/10.1093/jisesa/iead092.

Renkema JM, Iglesias LE, Bonneau P, Liburd OE. 2018. Trapping system comparisons for and factors affecting populations of Drosophila suzukii and Zaprionus indianus in winter-grown strawberry. Pest Management Science. 74(9):2076–2088. doi:10.1002/ps.4904. [accessed 2025 Mar 29]. https://onlinelibrary.wiley.com/doi/abs/10.1002/ps.4904.

Renkema JM, Miller M, Fraser H, Légaré J-P, Hallett RH. 2013. First records of *Zaprionus indianus* Gupta (Diptera: Drosophilidae) from commercial fruit fields in Ontario and Quebec, Canada. The Journal of the Entomological Society of Ontario. 144. [accessed 2021 Jan 25]. https://journal.lib.uoguelph.ca/index.php/eso/article/view/3745.

Ricciardi A. 2007. Are modern biological invasions an unprecedented form of global change? Conserv Biol. 21(2):329–336. doi:10.1111/j.1523-1739.2006.00615.x.

Roman G, Meller V, Wu KH, Davis RL. 1998. The opt1 gene ofDrosophila melanogaster encodes a proton-dependent dipeptide transporter. American Journal of Physiology-Cell Physiology. 275(3):C857–C869. doi:10.1152/ajpcell.1998.275.3.C857. [accessed 2024 Sep 4]. https://journals.physiology.org/doi/full/10.1152/ajpcell.1998.275.3.C857.

Roque F, Matavelli C, Lopes PHS, Machida WS, Von Zuben CJ, Tidon R. 2017. Brazilian Fig Plantations Are Dominated by Widely Distributed Drosophilid Species (Diptera: Drosophilidae). Annals of the Entomological Society of America. 110(6):521–527. doi:10.1093/aesa/sax044.

Sabeti PC, Reich DE, Higgins JM, Levine HZP, Richter DJ, Schaffner SF, Gabriel SB, Platko JV, Patterson NJ, McDonald GJ, et al. 2002. Detecting recent positive selection in the human genome from haplotype structure. Nature. 419(6909):832–837. doi:10.1038/nature01140. [accessed 2025 May 28]. https://www.nature.com/articles/nature01140.

Sabeti PC, Varilly P, Fry B, Lohmueller J, Hostetter E, Cotsapas C, Xie X, Byrne EH, McCarroll SA, Gaudet R, et al. 2007. Genome-wide detection and characterization of positive selection in human populations. Nature. 449(7164):913–918. doi:10.1038/nature06250. [accessed 2021 Feb 8]. https://www.ncbi.nlm.nih.gov/pmc/articles/PMC2687721/.

Sardain A, Sardain E, Leung B. 2019. Global forecasts of shipping traffic and biological invasions to 2050. Nature Sustainability. 2(4):274–282. doi:10.1038/s41893-019-0245-y.

Schluter D, Marchinko KB, Arnegard ME, Zhang H, Brady SD, Jones FC, Bell MA, Kingsley DM. 2021. Fitness maps to a large-effect locus in introduced stickleback populations. PNAS. 118(3). doi:10.1073/pnas.1914889118. [accessed 2021 Jan 26]. https://www.pnas.org/content/118/3/e1914889118.

Schmidt JM, Good RT, Appleton B, Sherrard J, Raymant GC, Bogwitz MR, Martin J, Daborn PJ, Goddard ME, Batterham P, et al. 2010. Copy Number Variation and Transposable Elements Feature in Recent, Ongoing Adaptation at the Cyp6g1 Locus. PLOS Genetics. 6(6):e1000998. doi:10.1371/journal.pgen.1000998. [accessed 2025 May 27]. https://journals.plos.org/plosgenetics/article?id=10.1371/journal.pgen.1000998.

Scott JG, Liu N, Wen Z. 1998. Insect cytochromes P450: diversity, insecticide resistance and tolerance to plant toxins1. Comparative Biochemistry and Physiology Part C: Pharmacology, Toxicology and Endocrinology. 121(1):147–155. doi:10.1016/S0742-8413(98)10035-X. [accessed 2025 May 29]. https://www.sciencedirect.com/science/article/pii/S074284139810035X.

Seebens H, Blackburn TM, Dyer EE, Genovesi P, Hulme PE, Jeschke JM, Pagad S, Pyšek P, Winter M, Arianoutsou M, et al. 2017. No saturation in the accumulation of alien species worldwide. Nature Communications. 8(1):14435. doi:10.1038/ncomms14435.

Seebens H, Essl F, Dawson W, Fuentes N, Moser D, Pergl J, Pyšek P, Kleunen M van, Weber E, Winter M, et al. 2015. Global trade will accelerate plant invasions in emerging economies under climate change. Global Change Biology. 21(11):4128–4140. 10.1111/gcb.13021.

Shi L, Li W, Dong Y, Shi Y, Zhou Y, Liao X. 2021. NADPH-cytochrome P450 reductase potentially involved in indoxacarb resistance in *Spodoptera litura*. Pesticide Biochemistry and Physiology. 173:104775. doi:10.1016/j.pestbp.2021.104775. [accessed 2025 May 29]. https://www.sciencedirect.com/science/article/pii/S0048357521000067.

Siddiqui JA, Fan R, Naz H, Bamisile BS, Hafeez M, Ghani MI, Wei Y, Xu Y, Chen X. 2023. Insights into insecticide-resistance mechanisms in invasive species: Challenges and control strategies. Front Physiol. 13. doi:10.3389/fphys.2022.1112278. [accessed 2025 Jun 4]. https://www.frontiersin.org/journals/physiology/articles/10.3389/fphys.2022.1112278/full.

da Silva VH, Laine VN, Bosse M, Spurgin LG, Derks MFL, van Oers K, Dibbits B, Slate J, Crooijmans RPMA, Visser ME, et al. 2019. The Genomic Complexity of a Large Inversion in Great Tits. Genome Biology and Evolution. 11(7):1870–1881. doi:10.1093/gbe/evz106. [accessed 2024 Jun 24]. https://doi.org/10.1093/gbe/evz106.

Simão FA, Waterhouse RM, Ioannidis P, Kriventseva EV, Zdobnov EM. 2015. BUSCO: assessing genome assembly and annotation completeness with single-copy orthologs. Bioinformatics. 31(19):3210–3212. doi:10.1093/bioinformatics/btv351. [accessed 2023 Jul 13]. https://doi.org/10.1093/bioinformatics/btv351.

Smit A, Hubley R, Green P. 2015. RepeatMasker Open-4.0. http://www.repeatmasker.org.

Soudi S, Crepeau M, Collier TC, Lee Y, Cornel AJ, Lanzaro GC. 2023. Genomic signatures of local adaptation in recent invasive Aedes aegypti populations in California. BMC Genomics. 24(1):311. doi:10.1186/s12864-023-09402-5. [accessed 2024 Jun 17]. https://doi.org/10.1186/s12864-023-09402-5.

Steenwyk JL, Rokas A. 2021. ggpubfigs: Colorblind-Friendly Color Palettes and ggplot2 Graphic System Extensions for Publication-Quality Scientific Figures. Microbiology Resource Announcements. 10(44): 10.1128/mra.00871-21. doi:10.1128/mra.00871-21. [accessed 2023 Jul 13]. https://journals.asm.org/doi/10.1128/MRA.00871-21.

Stern DB, Lee CE. 2020. Evolutionary origins of genomic adaptations in an invasive copepod. Nat Ecol Evol. 4(8):1084–1094. doi:10.1038/s41559-020-1201-y. [accessed 2024 Jun 18]. https://www.nature.com/articles/s41559-020-1201-y.

Stuart KC, Cardilini APA, Cassey P, Richardson MF, Sherwin WB, Rollins LA, Sherman CDH. 2021. Signatures of selection in a recent invasion reveal adaptive divergence in a highly vagile invasive species. Molecular Ecology. 30(6):1419–1434. 10.1111/mec.15601. [accessed 2021 Jan 5]. https://onlinelibrary.wiley.com/doi/abs/10.1111/mec.15601.

Sutherst RW, Constable F, Finlay KJ, Harrington R, Luck J, Zalucki MP. 2011. Adapting to crop pest and pathogen risks under a changing climate. WIREs Climate Change. 2(2):220–237. 10.1002/wcc.102.

Suvorov A, Kim BY, Wang J, Armstrong EE, Peede D, D’Agostino ERR, Price DK, Waddell PJ, Lang M, Courtier-Orgogozo V, et al. 2022. Widespread introgression across a phylogeny of 155 *Drosophila* genomes. Current Biology. 32(1):111–123.e5. doi:10.1016/j.cub.2021.10.052. [accessed 2024 Jun 21]. https://www.sciencedirect.com/science/article/pii/S0960982221014962.

Tait G, Mermer S, Stockton D, Lee J, Avosani S, Abrieux A, Anfora G, Beers E, Biondi A, Burrack H, et al. 2021. Drosophila suzukii (Diptera: Drosophilidae): A Decade of Research Towards a Sustainable Integrated Pest Management Program. Journal of Economic Entomology. 114(5):1950–1974. doi:10.1093/jee/toab158. [accessed 2025 Jun 9]. https://doi.org/10.1093/jee/toab158.

Tepolt CK, Grosholz ED, de Rivera CE, Ruiz GM. 2022. Balanced polymorphism fuels rapid selection in an invasive crab despite high gene flow and low genetic diversity. Molecular Ecology. 31(1):55–69. doi:10.1111/mec.16143. [accessed 2024 Jun 17]. https://onlinelibrary.wiley.com/doi/abs/10.1111/mec.16143.

Tepolt CK, Palumbi SR. 2020. Rapid Adaptation to Temperature via a Potential Genomic Island of Divergence in the Invasive Green Crab, Carcinus maenas. Front Ecol Evol. 8. doi:10.3389/fevo.2020.580701. [accessed 2024 Jun 20]. https://www.frontiersin.org/articles/10.3389/fevo.2020.580701.

Terhorst J, Kamm JA, Song YS. 2017. Robust and scalable inference of population history from hundreds of unphased whole-genomes. Nat Genet. 49(2):303–309. doi:10.1038/ng.3748. [accessed 2023 Jul 13]. https://www.ncbi.nlm.nih.gov/pmc/articles/PMC5470542/.

Thompson MJ, Jiggins CD. 2014. Supergenes and their role in evolution. Heredity. 113(1):1–8. doi:10.1038/hdy.2014.20. [accessed 2015 Aug 11]. http://www.nature.com/hdy/journal/v113/n1/full/hdy201420a.html.

Thornton T, Tang H, Hoffmann TJ, Ochs-Balcom HM, Caan BJ, Risch N. 2012. Estimating kinship in admixed populations. Am J Hum Genet. 91(1):122–138. doi:10.1016/j.ajhg.2012.05.024.

Timmeren SV, Isaacs R. 2014. Drosophila suzukii in Michigan vineyards, and the first report of Zaprionus indianus from this region. Journal of Applied Entomology. 138(7):519–527. 10.1111/jen.12113.

Uller T, Leimu R. 2011. Founder events predict changes in genetic diversity during human-mediated range expansions. Global Change Biology. 17(11):3478–3485. doi:10.1111/j.1365-2486.2011.02509.x. [accessed 2024 Jun 21]. https://onlinelibrary.wiley.com/doi/abs/10.1111/j.1365-2486.2011.02509.x.

Vilela C. 1999. Is Zaprionus indianus Gupta, 1970 (Diptera, Drosophilidae) currently colonizing the Neotropical region? Drosophila Information Service. 82:37–39.

Voight BF, Kudaravalli S, Wen X, Pritchard JK. 2006. A Map of Recent Positive Selection in the Human Genome. PLOS Biology. 4(3):e72. doi:10.1371/journal.pbio.0040072. [accessed 2019 Apr 17]. https://journals.plos.org/plosbiology/article?id=10.1371/journal.pbio.0040072.

Wang K, Peng X, Zuo Y, Li Y, Chen M. 2016. Molecular Cloning, Expression Pattern and Polymorphisms of NADPH-Cytochrome P450 Reductase in the Bird Cherry-Oat Aphid Rhopalosiphum padi (L.). PLOS ONE. 11(4):e0154633. doi:10.1371/journal.pone.0154633. [accessed 2025 May 26]. https://journals.plos.org/plosone/article?id=10.1371/journal.pone.0154633.

Wang X, Chen X, Zhou T, Dai W, Zhang C. 2025. NADPH-cytochrome P450 reductase mediates resistance to neonicotinoid insecticides in *Bradysia odoriphaga*. Pesticide Biochemistry and Physiology. 211:106406. doi:10.1016/j.pestbp.2025.106406. [accessed 2025 May 23]. https://www.sciencedirect.com/science/article/pii/S0048357525001191.

Whiting JR, Paris JR, Zee MJ van der, Parsons PJ, Weigel D, Fraser BA. 2021. Drainage-structuring of ancestral variation and a common functional pathway shape limited genomic convergence in natural high- and low-predation guppies. PLOS Genetics. 17(5):e1009566. doi:10.1371/journal.pgen.1009566. [accessed 2025 May 27]. https://journals.plos.org/plosgenetics/article?id=10.1371/journal.pgen.1009566.

Wickham H. 2016. ggplot2: Elegant Graphics for Data Analysis.

Willbrand B, Pfeiffer D, Leblanc L, Yassin A. 2018. First Report of African Fig Fly, Zaprionus indianus Gupta (Diptera: Drosophilidae), on the Island of Maui, Hawaii, USA, in 2017 and Potential Impacts to the Hawaiian Entomofauna. Proceedings of the Hawaiian Entomological Society. 50:55–65.

Yassin A, Capy P, Madi-Ravazzi L, Ogereau D, David JR. 2008. DNA barcode discovers two cryptic species and two geographical radiations in the invasive drosophilid Zaprionus indianus. Molecular Ecology Resources. 8(3):491–501. doi:10.1111/j.1471-8286.2007.02020.x. [accessed 2019 Aug 15]. https://onlinelibrary.wiley.com/doi/abs/10.1111/j.1471-8286.2007.02020.x.

Yassin A, David J. 2010. Revision of the Afrotropical species of Zaprionus (Diptera, Drosophilidae), with descriptions of two new species and notes on the internal reproductive structures and immature stages. ZooKeys. 51:33–72. doi:10.3897/zookeys.51.380.

Zanuncio-Junior JS, Fornazier MJ, Andreazza F, Culik MP, Mendonça L de P, Oliveira EE, Martins D dos S, Fornazier ML, Costa H, Ventura JA. 2018. Spread of Two Invasive Flies (Diptera: Drosophilidae) Infesting Commercial Fruits in Southeastern Brazil. flen. 101(3):522–525. doi:10.1653/024.101.0328.

Zheng X, Levine D, Shen J, Gogarten SM, Laurie C, Weir BS. 2012. A high-performance computing toolset for relatedness and principal component analysis of SNP data. Bioinformatics. 28(24):3326–3328. doi:10.1093/bioinformatics/bts606.

