## Supplementary Material for "Limited population structure but signals of recent selection in introduced African Fig Fly *(Zaprionus indianus*) in North America"

*PA Erickson et al*

**Table S1. Samples from previous work included in this study**

| Location | Year | Number Sequenced | Number after filtering | Source |
|---|---|---|---|---|
| Sao Tome | 2015 | 6 | 6 | Comeault 2020 |
| Senegal (Desert) | 2015 | 7 | 7 | Comeault 2020 |
| Senegal (Forest) | 2015 | 7 | 7 | Comeault 2020 |
| Zambia | 2015 | 6 | 6 | Comeault 2020 |
| Kenya | 2015 | 7 | 7 | Comeault 2020 |
| Hawaii, US (HI) | 2015 | 4 | 4 | Comeault 2020 |
| North Carolina, US (NC) | 2015 | 6 | 6 | Comeault 2020 |
| Tennessee, US (TN) | 2015 | 4 | 4 | Comeault 2020 |
| Florida, US (FL) | 2016 | 2 | 2 | Comeault 2021 |
| New Jersey, US (NJ) | 2016 | 2 | 2 | Comeault 2021 |
| North Carolina, US (NC) | 2017 | 1 | 1 | Comeault 2021 |
| New York, US (NY) | 2017 | 2 | 2 | Comeault 2021 |
| Kenya | 2018 | 1 | 1 | Comeault 2021 |
| Colombia | 2018 | 4 | 4 | Comeault 2021 |
| North Carolina, US (NC) | 2018 | 6 | 6 | Comeault 2021 |
| Pennsylvania, US (PA) | 2018 | 2 | 2 | Comeault 2021 |

**Table S2: Muller elements in *Z. indianus*.** *D. melanogaster* similarities were identified using a dot-plot (Figure S5). Ananina *et al.* (2007) chromosome names were inferred based on relative chromosome sizes and presence of inversions.

| Muller element | *Z. indianus* chromosome (scaffold #) | Ananina *et al.* (2007) chromosome | *D. melanogaster* chromosome arm |
|---|---|---|---|
| A | 3 | X | X |
| B | 2 | V | 2L |
| C | 5 | II | 2R |
| D | 4 | III | 3L |
| E | 1 | IV | 3R |
| F | 8 | VI | 4 |

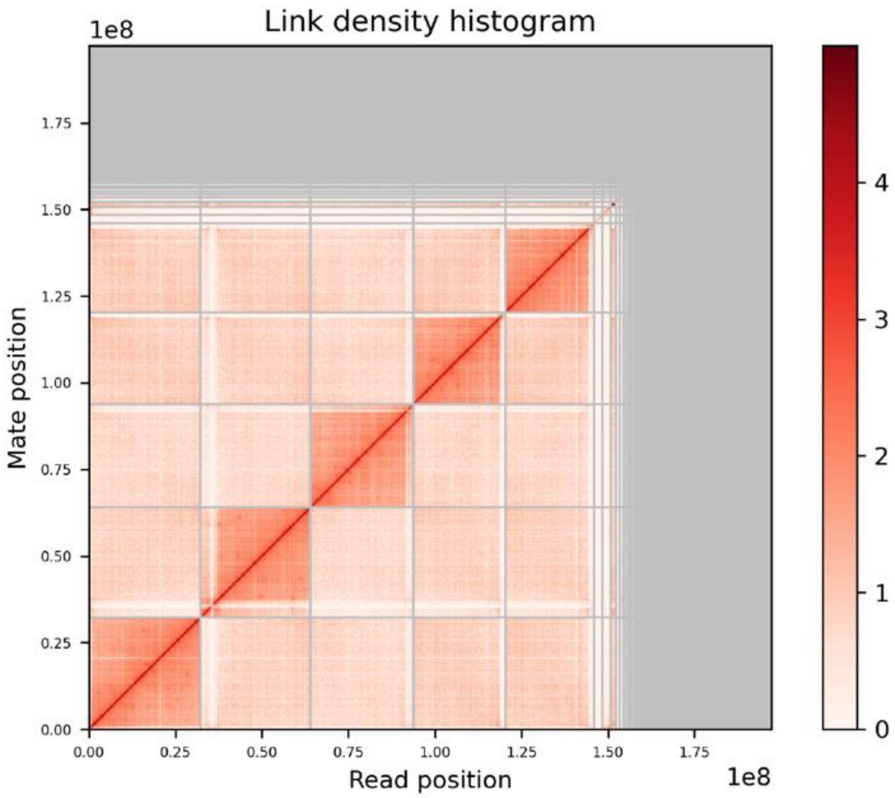


**Figure S1: Link density from Hi-C sequencing of *Z. indianus* genome illustrates physical contact between 5 major scaffolds.**

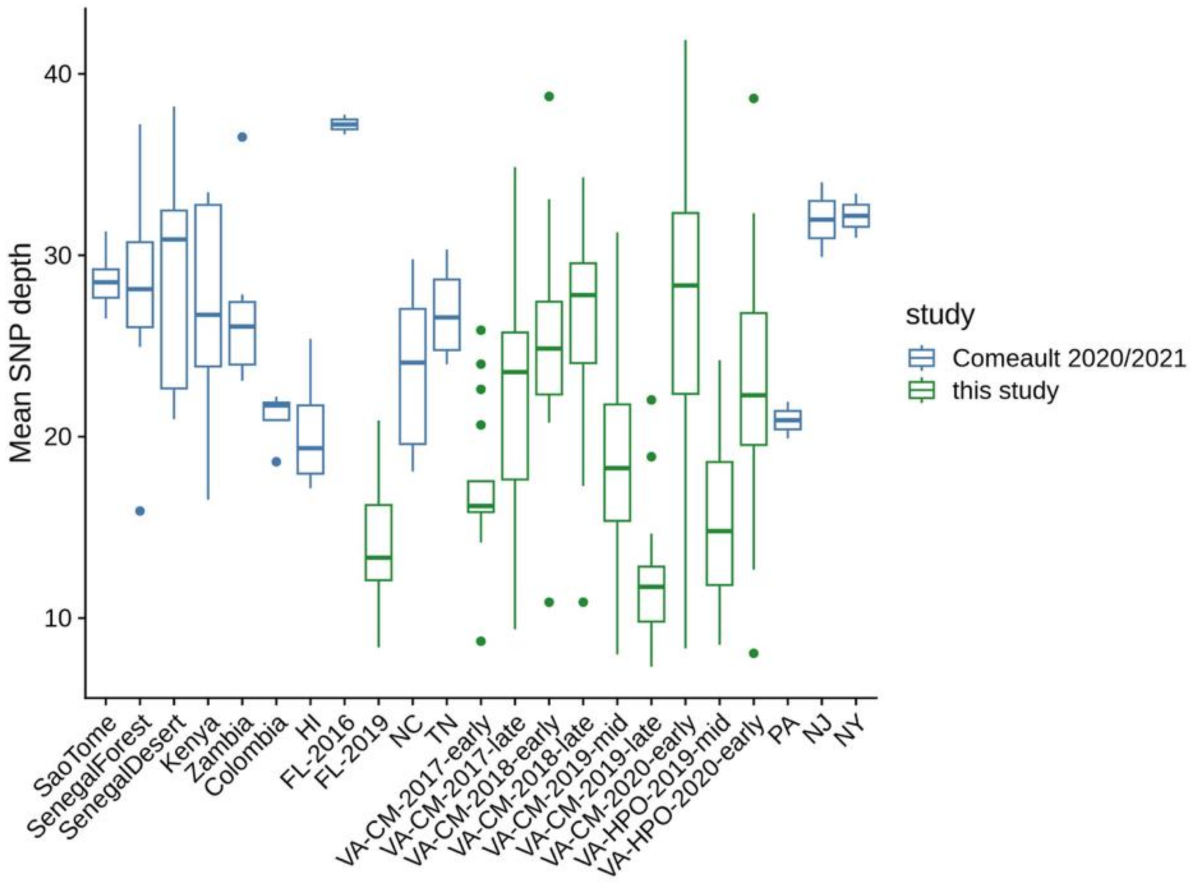


**Figure S2: Mean SNP depth by sampling location and date.** The mean depth was calculated across all SNPs (~5.18M sites) for each individual sample using *vcftools*. Samples are color coded based on whether they were previously published (blue; Comeault et al 2020, 2021) or sequenced for this study (green).

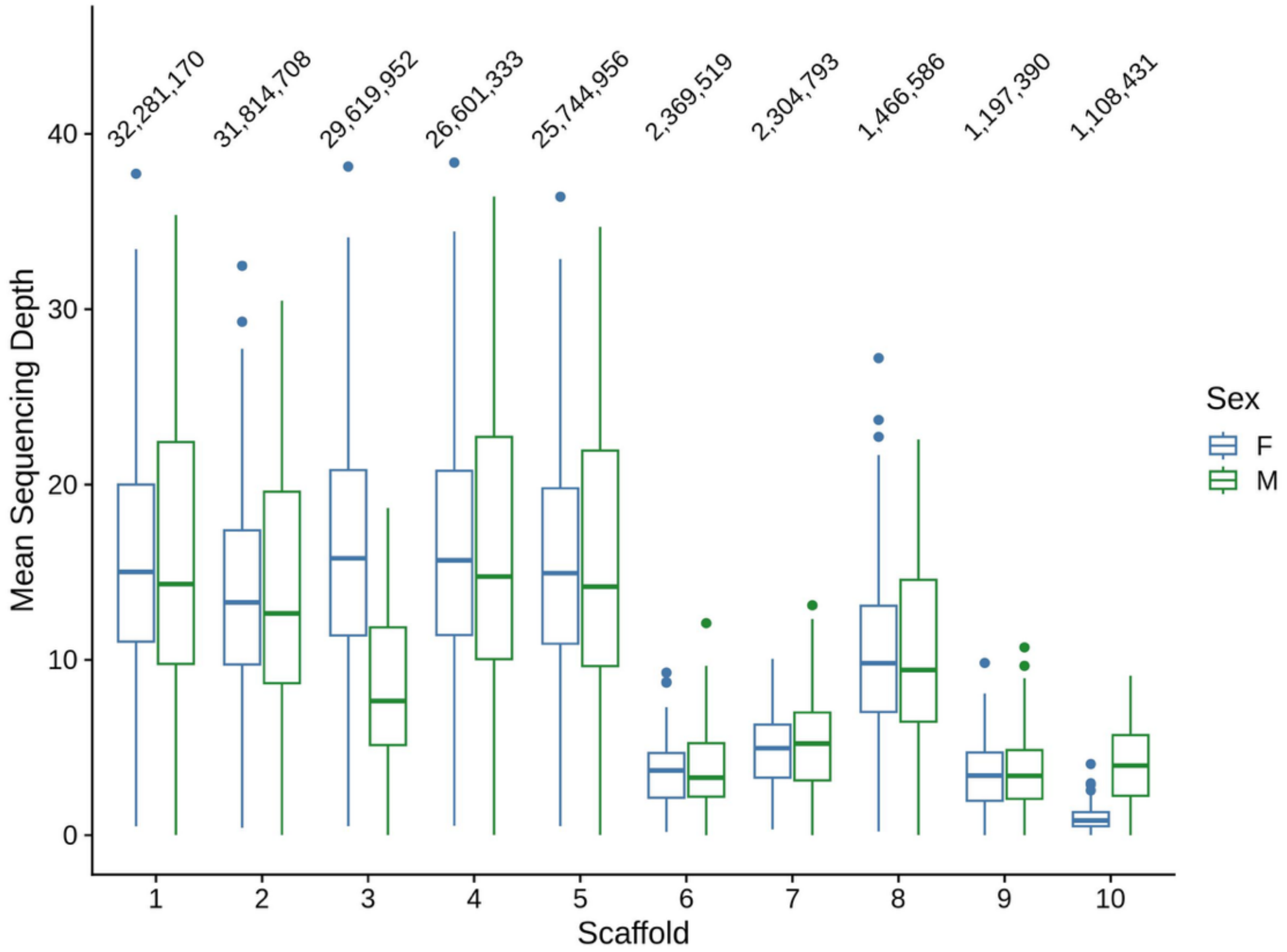


**Figure S3: Mean sequencing depth of samples for the ten largest scaffolds for samples with known sexes.** Each boxplot shows the distribution of sequencing coverage for n=286 samples. The total length of the scaffold in base pairs is given above the boxplot and sexes are shown separately. Scaffold 3 is the X chromosome.

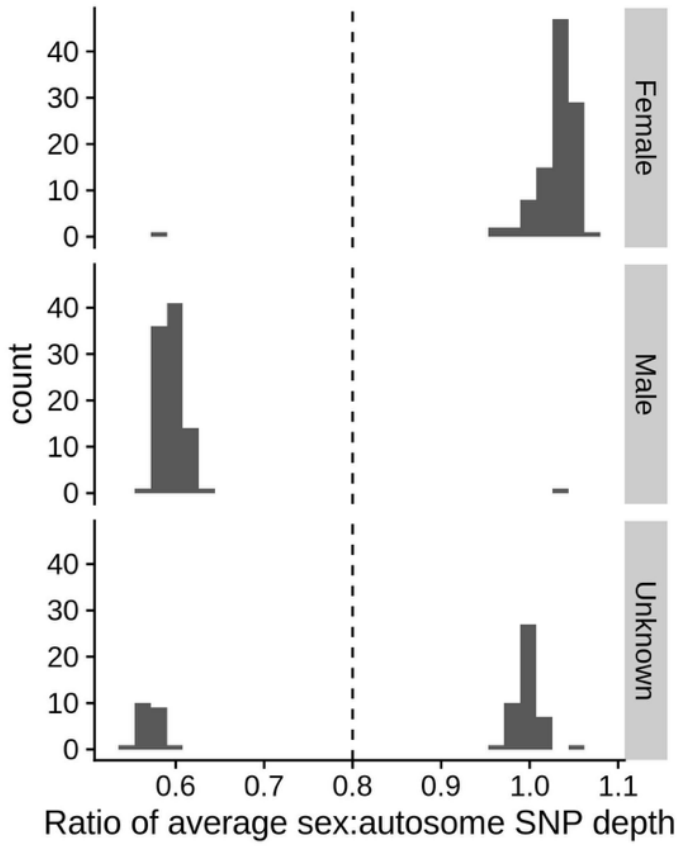


**Figure S4: Ratio of sex chromosome (scaffold 3) to autosomal (scaffolds 1, 2, 4, and 5) SNP depths based on sex of flies**. Sequencing depth was calculated for ~4.31M autosomal SNPs and ~860K X-chromosome SNPs in 105 female, 94 male, and 67 unknown sex samples. The dashed vertical line shows the cutoff used for assigning males (< 0.8) and females (>0.8) for unknown individuals. Two individuals that were likely misassigned sex at the time of DNA extraction were corrected based on these cutoffs.

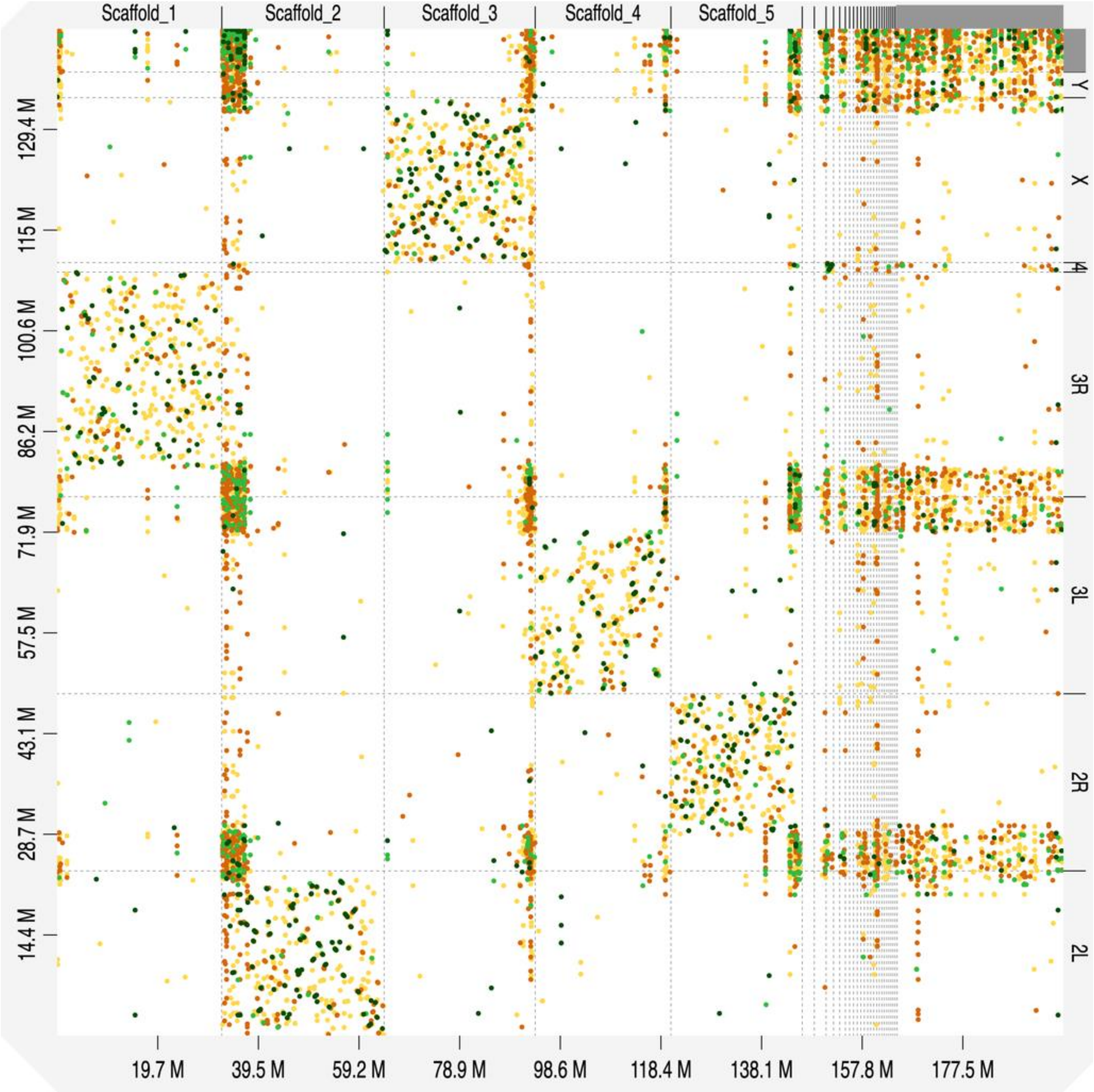


**Figure S5: Dot-plot comparison of *Z. indianus* and *D. melanogaster* genomes to identify Muller elements.** *Z. indianus* scaffolds are shown on the horizontal axis and *D. melanogaster* chromosomes arms are shown on the vertical axis. See Table S2 for Muller element assignments.

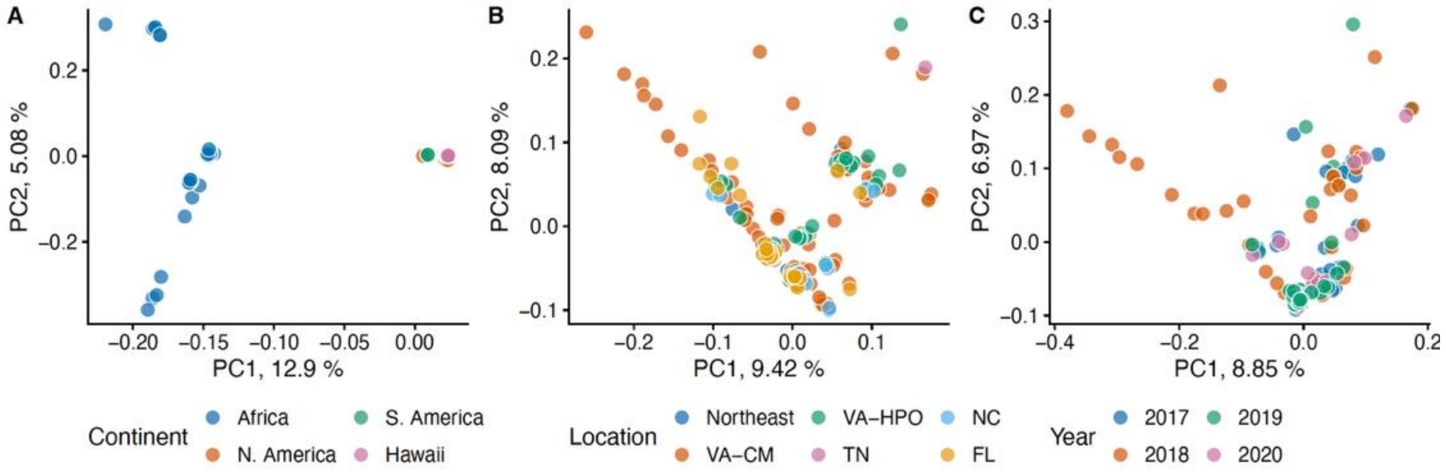


**Figure S6. Principal components analysis using autosomal SNPs.** Groupings are same as in Figure 1: A) all samples colored by continent, B) North American samples colored by location C) Carter Mountain, Virginia samples colored by year. PCs were recalculated for the subset of data included in each panel using LD-pruned SNPs with a minor allele count of at least 3. Regions potentially containing inversions were included in the analysis.

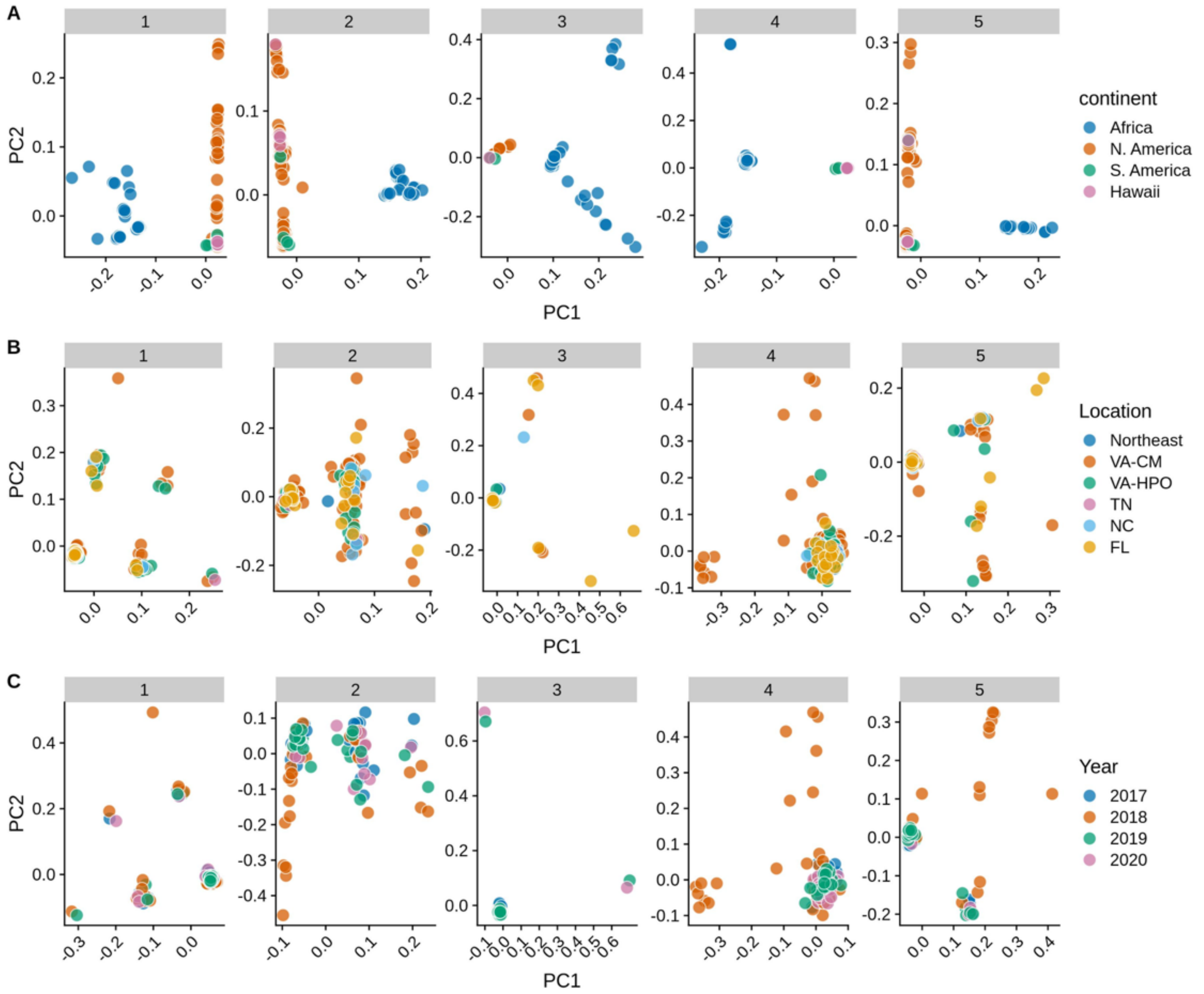


**Figure S7: Principal components analysis by chromosome.** Each facet represents PCA conducted using LD-pruned SNPs from one chromosome. Groupings are same as in Figure 1: A) all samples colored by continent, B) North American samples colored by location C) Carter Mountain, Virginia samples colored by year. Only females were used for the analysis of chromosome 3 (X), resulting in fewer data points. Regions potentially containing inversions were included in the analysis.

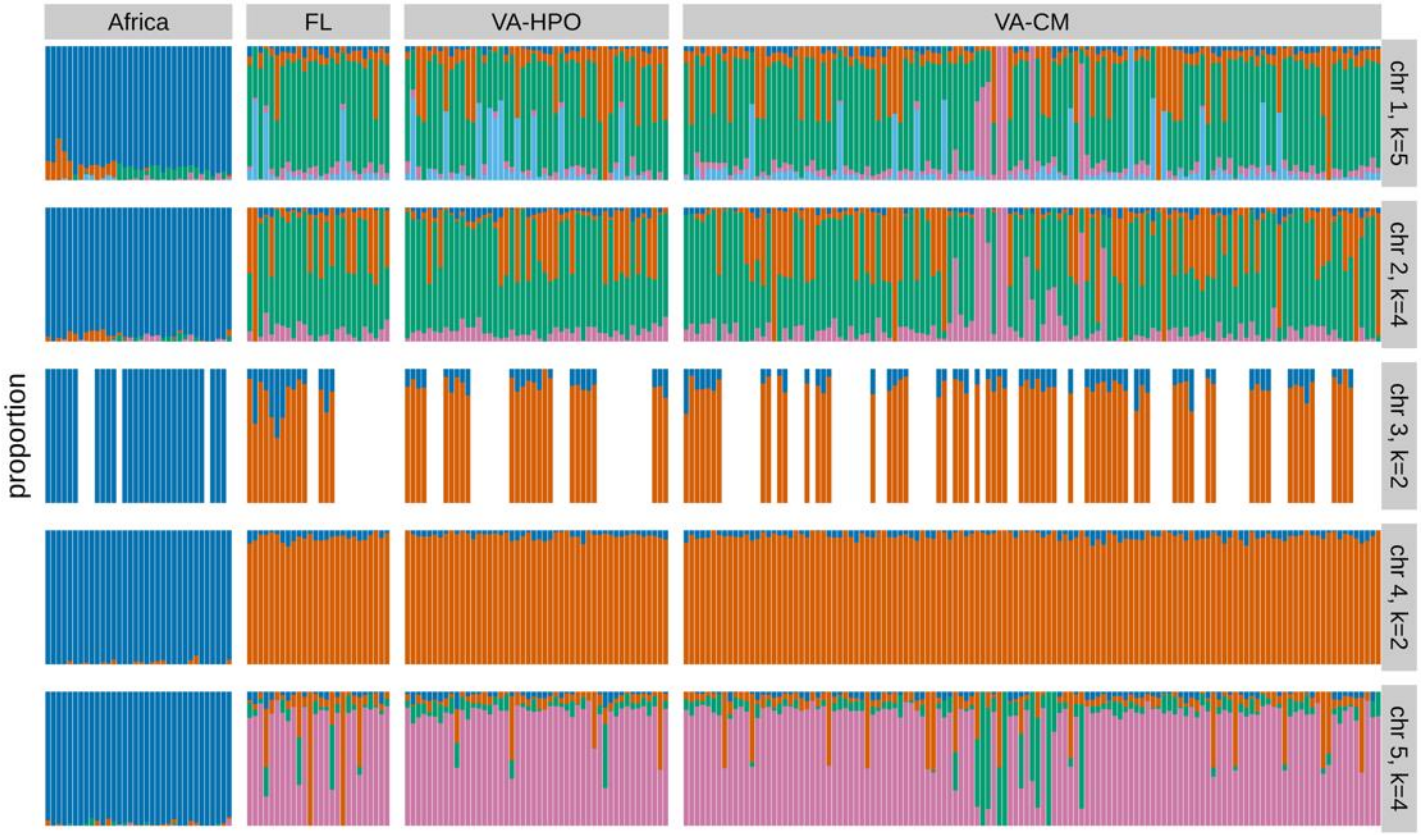


**Figure S8. Admixture analysis using individual chromosomes***.* Each column is an individual, each row represents data from a single chromosome, and colors represent assignment to distinct genetic clusters. The most likely number of genetic clusters for each chromosome (k) was obtained with cross-validation analysis and is shown at right. For chromosome 3, the X chromosome, only female flies were included, resulting in reduced sample size. African sequences represent five geographic locations and are taken from Comeault et al. (2020 & 2021). FL=Miami, Florida, VA-HPO = Hanover Peach Orchard, Virginia, VA-CM = Carter Mountain, Virginia.

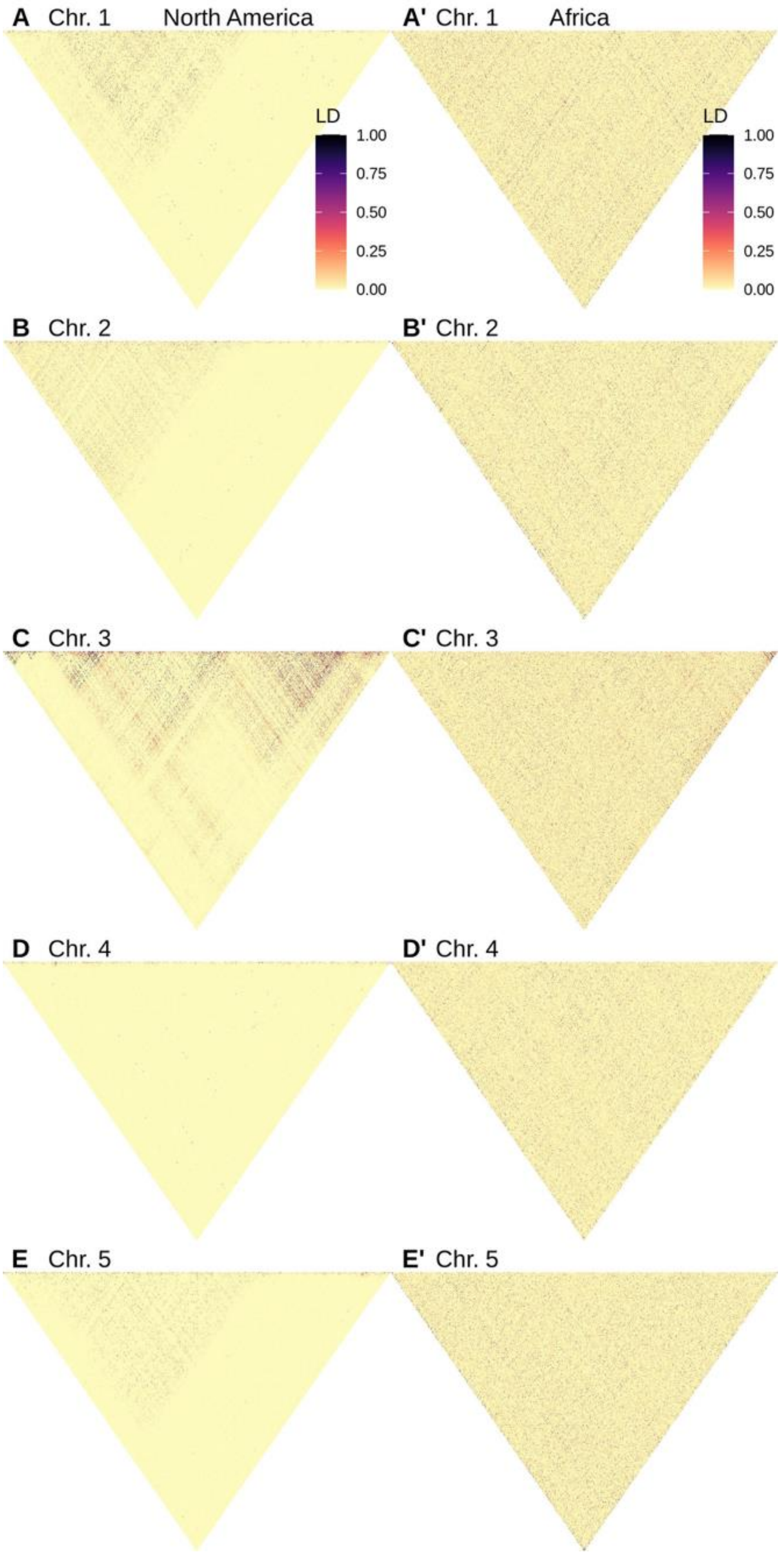


**Figure S9: Patterns of linkage disequilibrium on each chromosome indicate potential structural variation on all chromosomes except chromosome 4.** For each chromosome, 4000 random SNPs with no missing data were sampled. Panels A-E show $R^2$ in all North American samples (n=224 individuals; n=120 females for chromosome 3), and panels A'-E' show linkage patterns in Africa (n=34 individuals; n=28 females for chromosome 3).

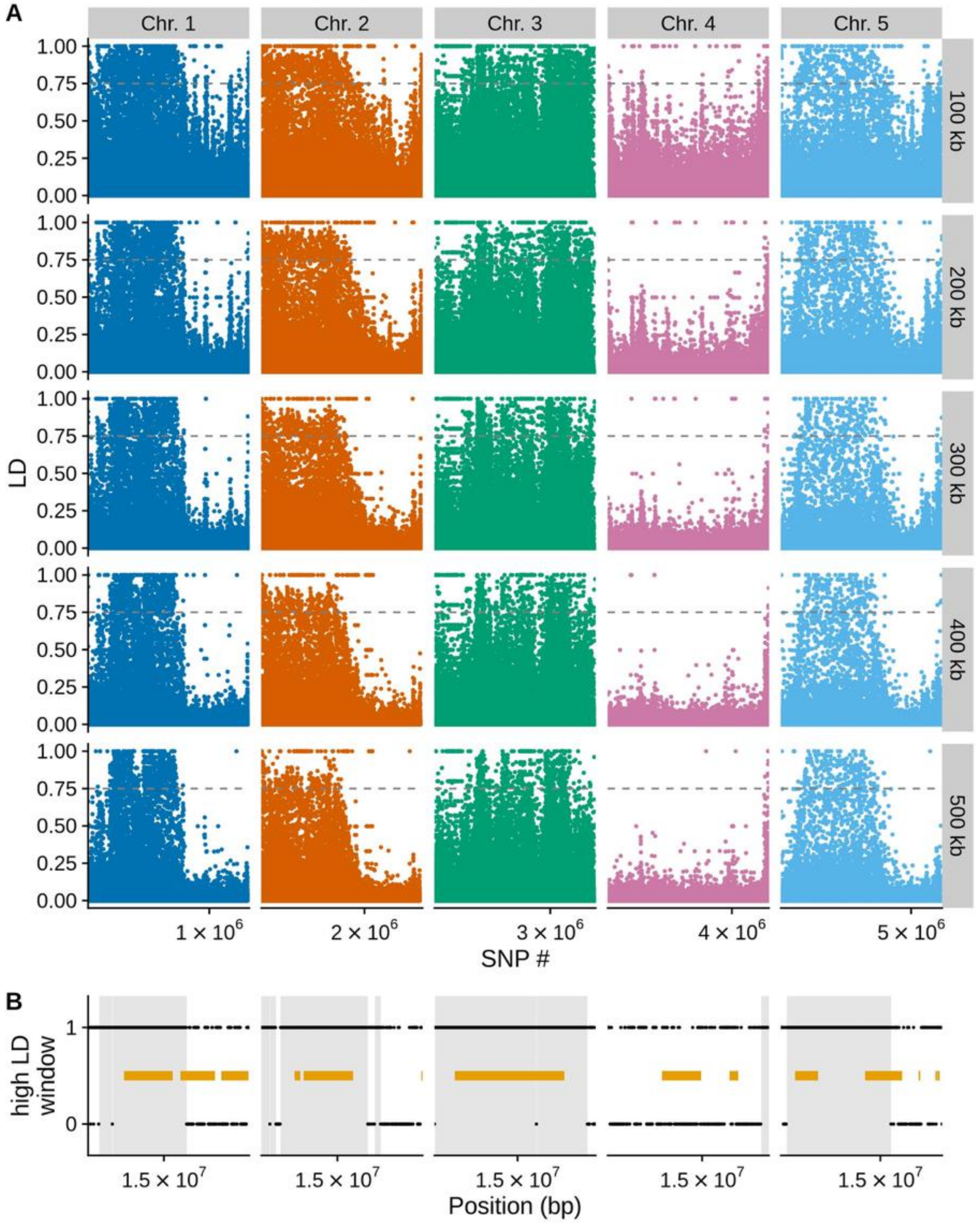


**Figure S10. Long-distance LD indicates presence of inversions in North American *Z. indianus*.** (A) For each segregating SNP, we randomly chose five other SNPs that were ~100, 200, 300, 400, and 500 kb away from the focal SNP and then calculated LD ($R^2$) between the two SNPs. LD is plotted for each focal SNP, with data divided into rows indicating each distance and columns indicating chromosomes. (B) We divided the genome into 100 kb non-overlapping windows and for each window determined if any SNPs in that window had long-distance LD of > 0.75 (high LD window = 1). Windows lacking high LD SNPs were assigned a value of zero. We defined inversions as any consecutive runs of high-LD windows that were 1 Mb or longer. Shaded regions show regions of each chromosome that were called as potential inversions and masked for population genomic analysis. Yellow rectangles show putative inversions > 100 kb as called by *smoove*; in general, *smoove* did not identify the same inversion boundaries as the LD analysis did.

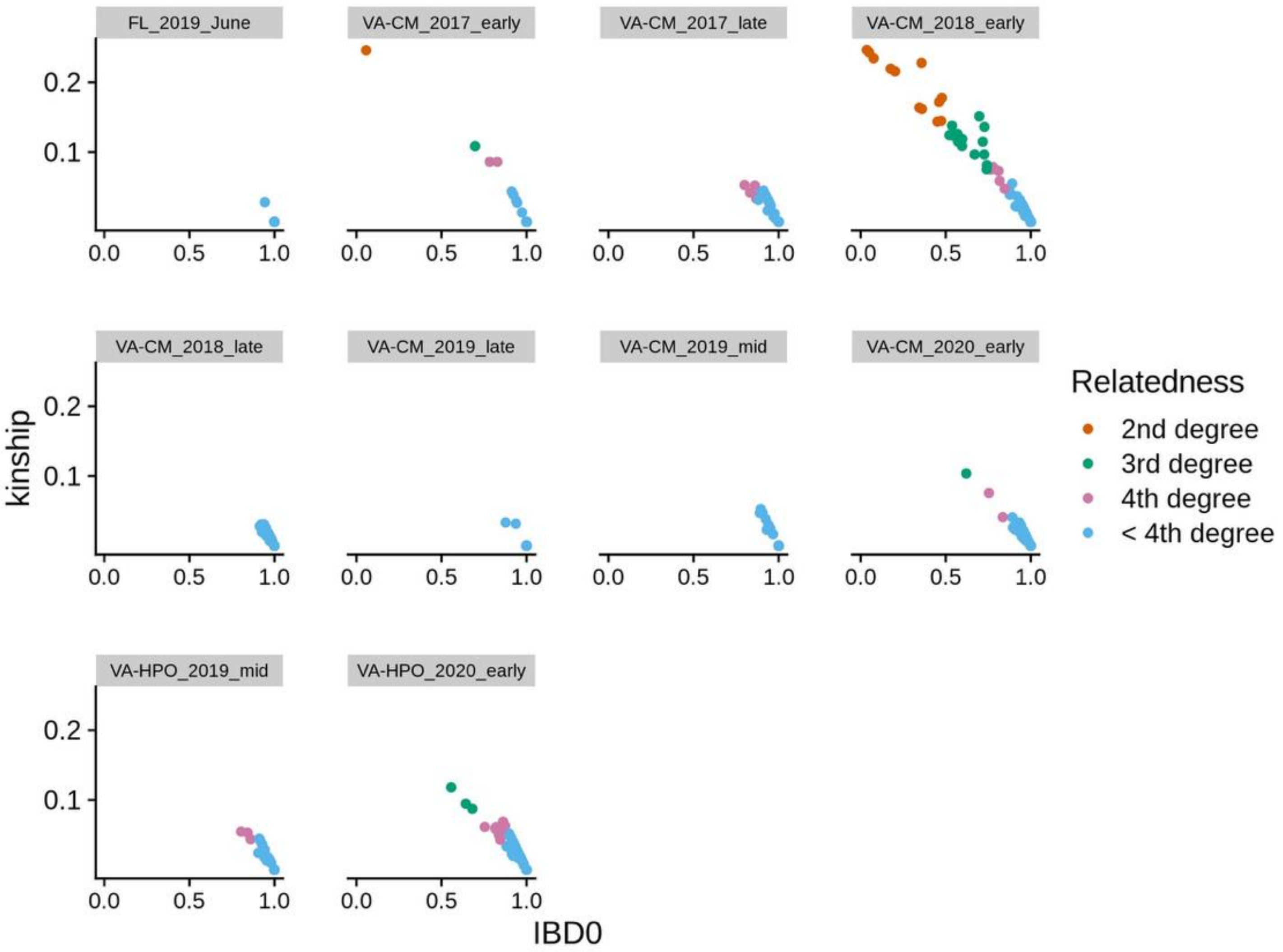


**Figure S11: Probability of zero identity by descent vs kinship for all pairs of individuals from North American populations.** Data are faceted by collection location and date. CM=Carter Mountain, Virginia, HPO=Hanover Peach Orchard, Virginia; early/mid/late refers to the timing of the collection). Color coding assigns relatedness according to (Thornton *et al.* 2012).

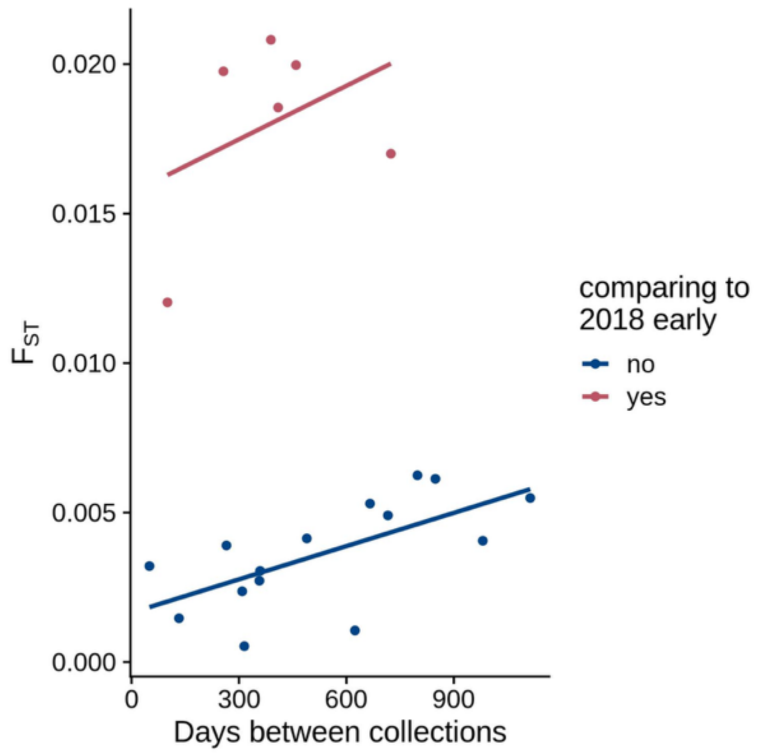


**Figure S12: $F_{ST}$ over time in the Carter Mountain, VA population.** Autosome-wide $F_{ST}$ was calculated for all pairwise combinations of sampling timepoints 2017-2020. Red points indicated comparisons that include the 2018 early sample (which contained many close relatives and is more genetically distant from other collections) and blue points indicate all other comparisons.

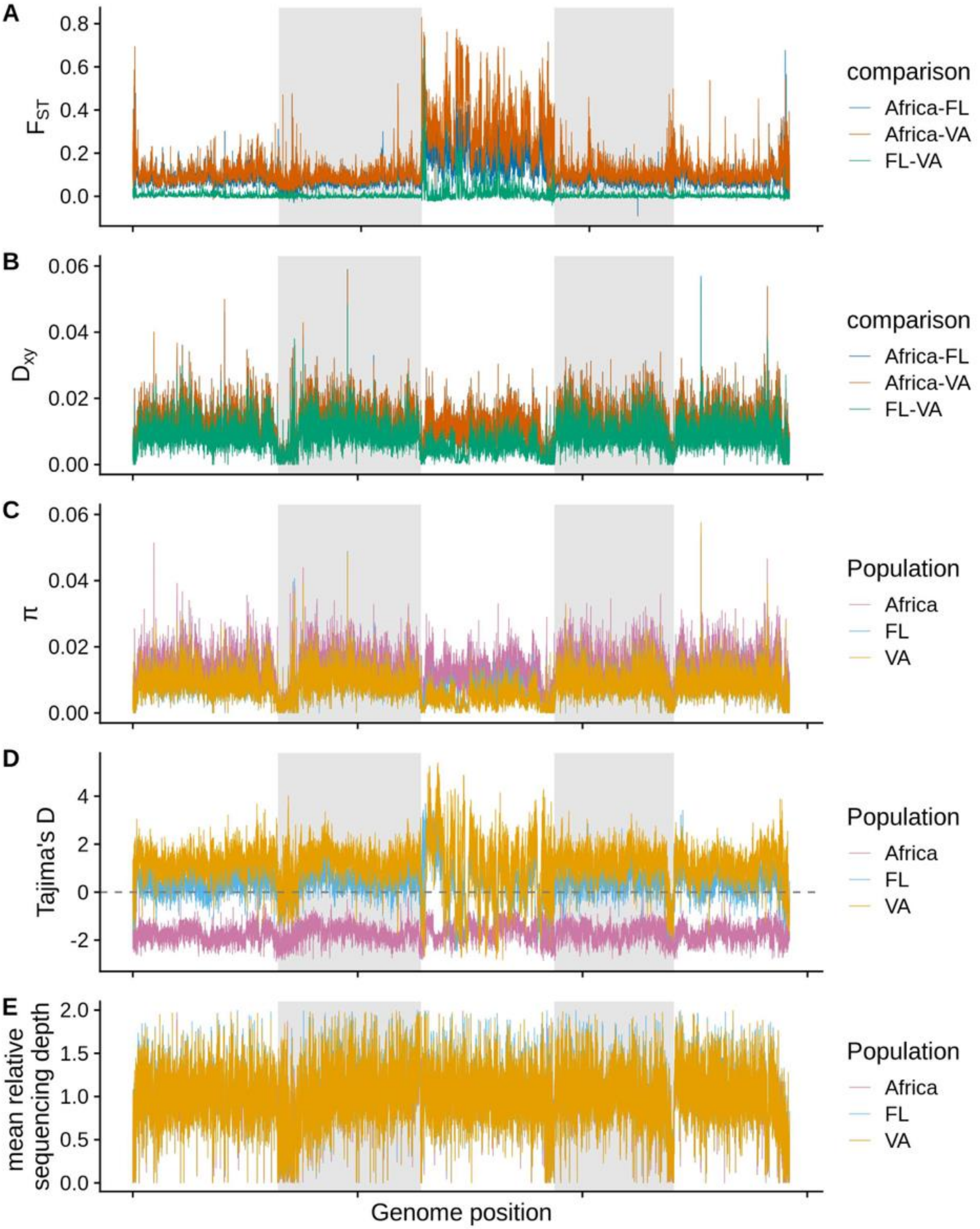


**Figure S13: Genome-wide population genetic and sequencing data.** All statistics were calculated in non-overlapping 5 kb windows, grouping flies into 3 major populations: Africa, Florida, and Virginia. Only females were used for calculations on the X chromosome. Alternate grey and white backgrounds indicate chromosome boundaries.

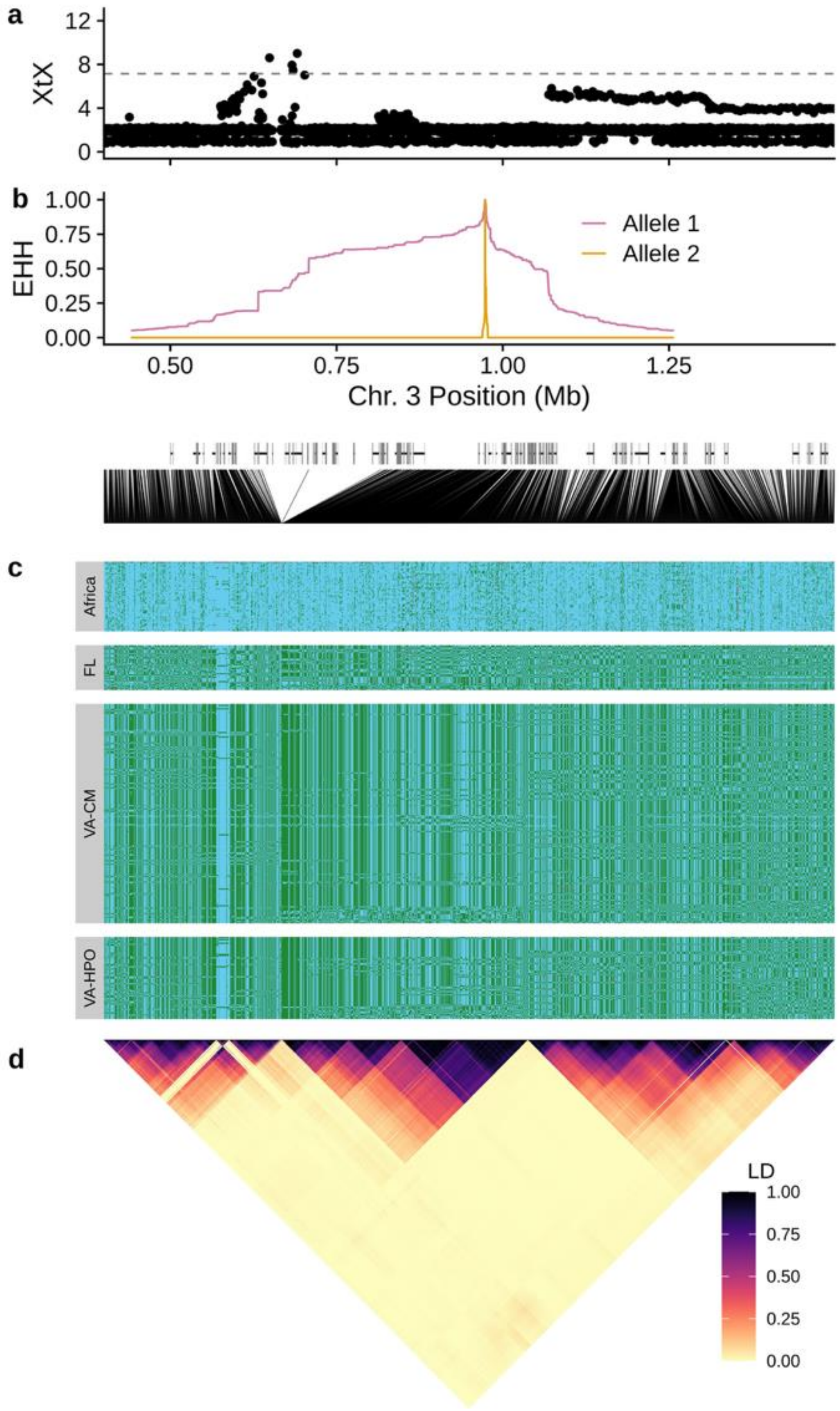


**Figure S14: A haplotype on the X chromosome with signals of selection and differentiation**. A) *BayPass* XtX score of individual SNPs comparing Florida and Virginia populations (see Figure 4). The dashed line indicates the 99.9% quantile of XtX scores for 10,000 simulated SNPs. B) Extended haplotype homozygosity plot for the SNP with highest IHS (Chromosome 3: 973,443) using genotypes from Virginia flies. Below the plot are locations of genes and lines to indicate the physical positions of SNPs show in panels c and d. C) Each row shows haplotypes of 2,004 SNPs for a single haploid chromosome phased with read-backed phasing. Blue indicates the allele more common in African populations and green is the other allele. Missing genotypes are shown in gray. D) LD ($R^2$) for these SNPs in North American flies, excluding Florida.

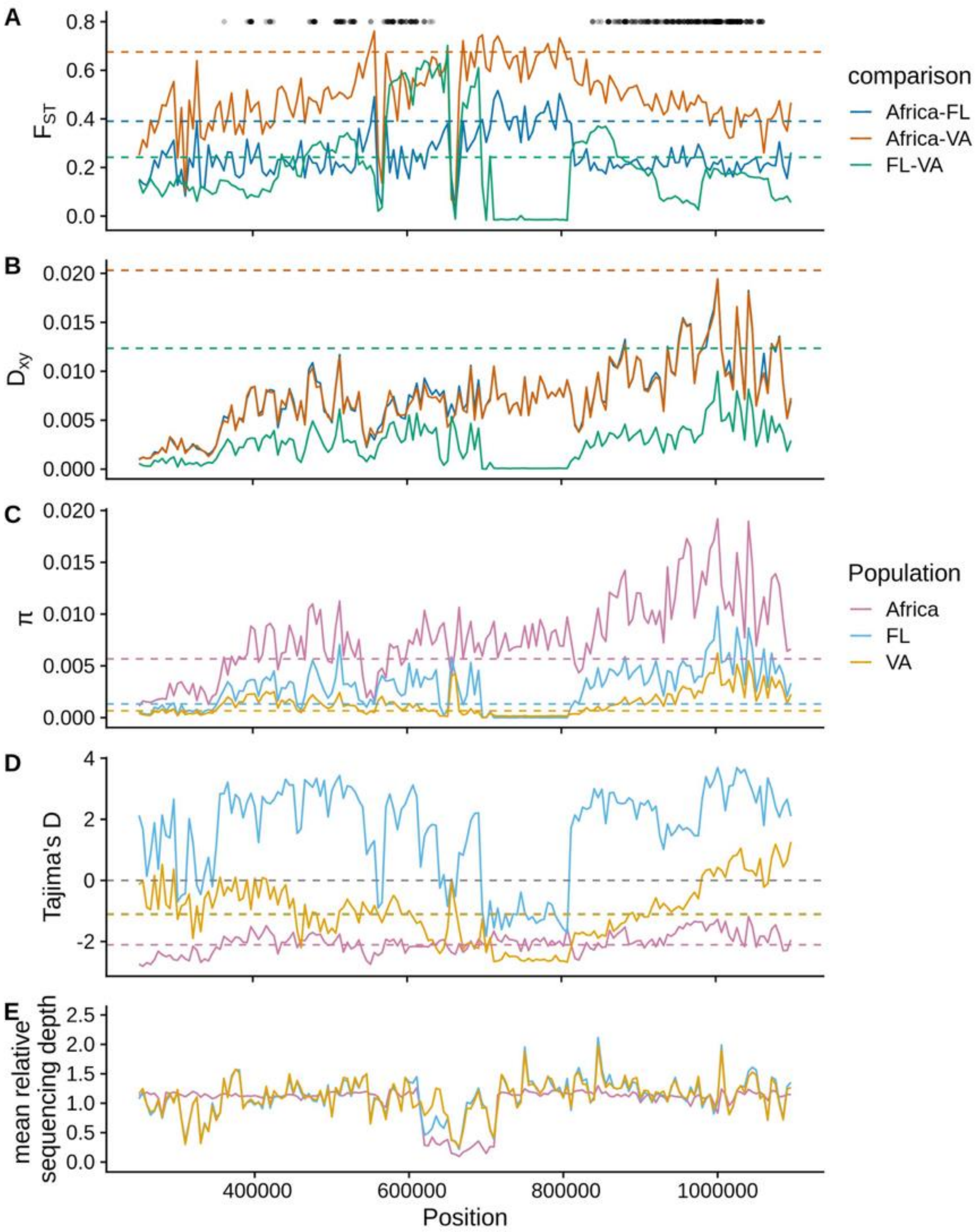


**Figure S15: Population genetic statistics for the region surrounding a potential selected haplotypes on chromosome 3 (250-1,1000 kb).** All statistics were calculated for 5 kb, non-overlapping windows. Black points at the top indicate the locations of 400 SNPs with IHS > 5. A) $F_{ST}$ comparing combinations of flies from Africa, Virginia (both focal orchards combined) and Florida. B) Absolute nucleotide divergence ($D_{xy}$) for the same comparisons. Dashed horizontal lines indicate 99% quantile for all X chromosome windows in A-B. C) Nucleotide diversity (π) for each population. D) Tajima's D for the three populations. Dashed horizontal lines indicate 1% quantile for all X chromosome windows in C-D. E) Average sequencing depth per window relative to the mean depth for the entire chromosome. Relative depths were averaged for all individuals in each population. See Figure S13 for whole-genome analysis of the same statistics.

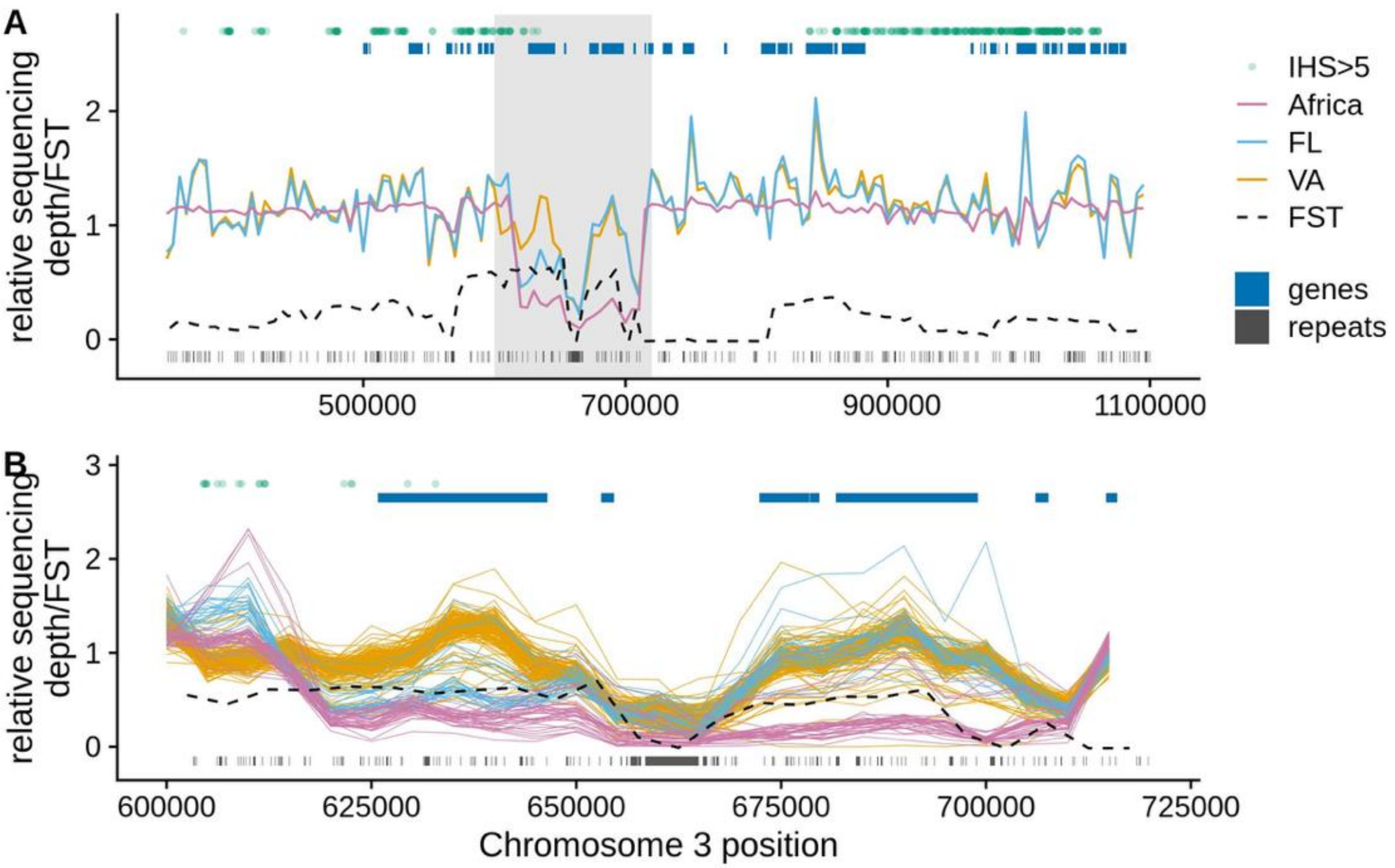


**Figure S16: Sequencing depth in the region of interest on chromosome 3.** A) Mean sequencing depth relative the rest of chromosome 3 of individuals from each population (pink, blue, and yellow lines). Mean relative depth was calculated in 5 kb non-overlapping windows and averaged across all individuals in each population. FST calculated in 5 kb windows is shown as the dashed black line. The locations of genes are shown as blue boxes at top, and repetitive regions are shown as gray boxes at the bottom. The location of the 400 SNPs with IHS > 5 are shown as green dots at the top. B) Zoom in on the gray region from (A). Each line shows relative sequencing depth for an individual, color coded by population.
